# Development of gemcitabine-modified miR-15a as a novel, multimodal treatment strategy to overcome 5-FU and oxaliplatin resistance in colorectal cancer

**DOI:** 10.64898/2026.04.25.720825

**Authors:** Anushka Ojha, Amartya Pal, Max Chao, Ramana V Davuluri, Jingfang Ju

## Abstract

**Background:** Resistance to 5-fluorouracil (5-FU)-based chemotherapy is a major clinical obstacle in colorectal cancer (CRC), highlighting the urgent need to overcome established resistance mechanisms. MicroRNA-based therapeutics have emerged as compelling candidates in this context, given their inherently pleiotropic mode of action; however, their clinical translation remains hindered by poor stability and suboptimal delivery.

**Methods:** To address these limitations, Gem-miR-15a, a unique gemcitabine-modified tumor-suppressor microRNA-15a was designed to synergistically integrate the tumor-suppressive activity of miR-15a with the chemotherapeutic potency of gemcitabine into a single molecular entity. Therapeutic efficacy of Gem-miR-15a was evaluated across a spectrum of preclinical models, including parental and drug-resistant CRC cell lines, 3D tumor spheroids, patient-derived organoids and *in vivo* metastatic models. Cell viability, apoptosis and cell cycle analyses were performed, along with RNA sequencing and protein validation. Statistical analyses were conducted using Student’s t-test or two-way ANOVA with mixed effects, and data were presented as mean ± SD.

**Results:** Gem-miR-15a exhibited potent anti-proliferative activity with IC_50_ values in the low nanomolar range, achieving ∼100-5000-fold greater potency relative to 5-FU and oxaliplatin. Importantly, it retained efficacy in both 5-FU- and oxaliplatin-resistant CRC models, effectively overcoming acquired chemoresistance. Mechanistically, Gem-miR-15a induced S-phase cell cycle arrest, eliminated the G2-phase cell population, and triggered apoptosis, accompanied by suppression of key oncogenic targets including WEE1, CHK1, YAP1 and BMI1. RNA-seq analysis further demonstrated modulation of pathways such as p53 signaling and reversal of resistance-associated gene expression, that were corroborated at the protein level. *In vivo*, Gem-miR-15a significantly reduced tumor growth at a dose ∼12-fold lower than gemcitabine, with no observable toxicity.

**Conclusion:** Gem-miR-15a represents a potent, multi-targeted therapeutic strategy capable of overcoming chemoresistance in CRC. Its enhanced stability, effective delivery and robust efficacy across resistant models and a favorable safety profile highlight its strong potential for clinical translation.

**Graphical Abstract:** 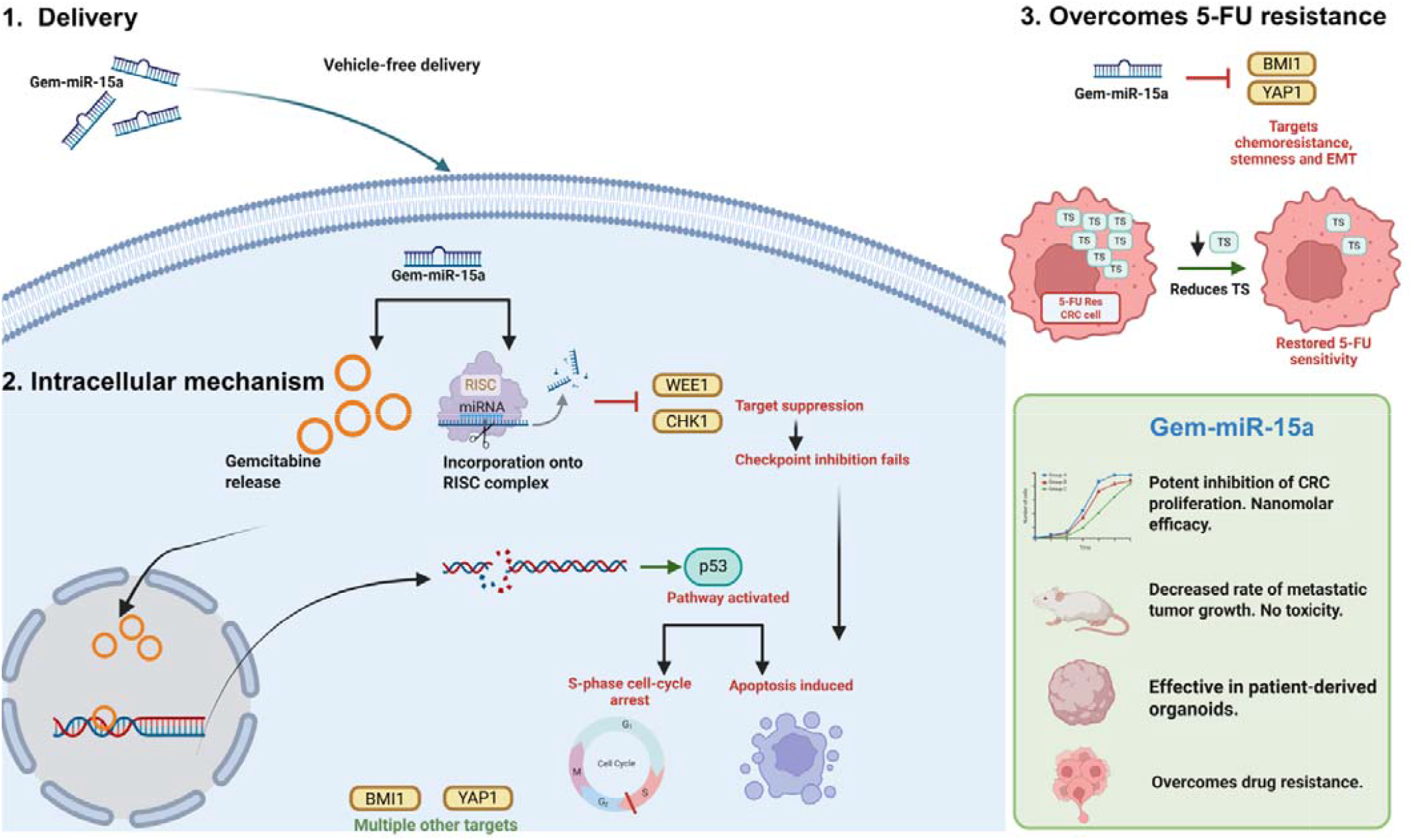

## Background

Colorectal cancer (CRC) is the second leading cause of cancer-related deaths in the United States, with an estimated 150,000+ new cases annually (1, 2). While overall incidence has dropped due to screening, cases are rising by ∼3% annually in adults under 50, with nearly half of all new cases now occurring under 65 (3). Despite improved screening strategies, which have improved the five-year survival rate for CRC to 64%, this figure drops to a dismal 14% for metastatic CRC (4). Treatment options in CRC primarily include surgery, chemotherapy and targeted therapies.

Treatment regimen is generally determined depending on disease stage, with the first-line therapy typically consisting of 5-FU-based chemotherapy in combination with leucovorin and oxaliplatin (Ox) (FOLFOX) or capecitabine and oxaliplatin (CAPOX). In advanced stage disease, these regimens are often combined with targeted therapies such as EGFR inhibitors (cetuximab) or VEGF inhibitors (bevacizumab) (5, 6). These combination strategies have improved clinical outcomes, leveraging the complementary mechanisms of 5-FU, a nucleoside analog that inhibits thymidylate synthesis (TS) and disrupts DNA/RNA synthesis and oxaliplatin, which forms platinum-DNA adducts that impede DNA replication (7, 8). Although immune checkpoint inhibitors (ICIs) have revolutionized treatment for mismatch repair-deficient metastatic CRC (∼4%), the majority of CRC patients do not benefit (9). Despite these therapeutic advances, resistance and relapse remain a major challenge in CRC, driven by tumor heterogeneity, dysregulation of oncogenic pathways, and the emergence of cancer stem cells (10). Advanced CRC shows a recurrence rate of ∼30% following primary treatment or surgery, with many patients eventually developing resistance to 5-FU and oxaliplatin, upon prolonged exposure (10, 11). Furthermore, 10-30 % of patients treated with fluoropyrimidines experience severe treatment-related toxicities (12).

Acquired resistance not only drives recurrence but also promotes pro-survival signaling and stemness (10). Elucidating the multifaceted mediators of resistance is therefore critical, with an urgent need to develop novel therapeutic strategies that overcome resistance by targeting regulatory networks simultaneously rather than individual targets (13). In this regard, miRNA-based therapeutics offer a compelling strategy to reprogram signaling networks and are being studied extensively as a promising avenue for improving treatment outcomes (14, 15). The awarding of the 2024 Nobel Prize in Physiology or Medicine for the discovery of miRNAs and their role in post-transcriptional gene regulation underscores their profound impact on biological systems, further driving the exploration of miRNAs as therapeutic agents (16). However, despite their considerable therapeutic potential, no FDA-approved miRNA-based therapeutic exists for cancer, primarily due to instability and delivery challenges, among other setbacks (15, 17).

MicroRNAs (miRNAs) are a family of short (∼20-22 nucleotide), non-coding RNAs that regulate a broad spectrum of biological processes through post-transcriptional gene silencing. They act by binding to the 3’ UTR of target mRNAs via a 6-8-mer ‘seed’ sequence located at their 5’ end (18-21). Remarkably, miRNA dysregulation has been implicated in a variety of diseases, including cancer (18, 21, 22). Due to their ability to bind to targets with incomplete complementarity, miRNAs can target multiple genes simultaneously, thereby modulating multiple pathways. This makes miRNAs a very attractive candidate for exploration and development of novel therapeutic strategies. In the cancer context, miRNA-based strategies have been explored through restoration of tumor-suppressor miRNAs or inhibition of oncogenic miRNAs (23-25).

Hsa-miR-15a (miR-15a) is one such tumor-suppressor miRNA, that has been studied extensively in the context of cancer and was in fact, the first miRNA to have been identified with an established role in cancer (26). Downregulation of miR-15a has been reported in multiple cancers like chronic lymphocytic leukemia, pituitary adenomas, and prostate carcinoma and is associated with cancer progression and metastases (27-29). Low miR-15a expression has previously been established to correlate with poor patient prognosis in many cancers, including pancreatic cancer, colorectal cancer and gastric cancers (30-32). The Cancer Genome Atlas (TCGA) analysis in advanced-stage CRC patients confirmed that low miR-15a expression correlates with poor prognosis (30, 33). Additionally, miR-15a targets key oncogenic targets such as WEE1, CHK1, YAP1 and BMI1 which play important roles in cell proliferation, self-renewal, metastases and chemoresistance, each independently validated as a promising therapeutic targets (30, 31, 34-38). miR-15a thereby suppresses progression and mitigates chemoresistance, by targeting multiple pathways simultaneously. Restoration of miR-15a levels or delivery of modified miR-15a mimics has demonstrated significant therapeutic potential (30, 31).

Gem-miR-15a was therefore designed as a gemcitabine-modified miR-15a (Gem-miR-15a), in which gemcitabine (Gem), a cytidine analog, was incorporated into the guide strand of hsa-miR-15a. Gem is widely used as a chemotherapeutic agent in multiple cancers and upon cellular uptake, is converted into its active form, gemcitabine triphosphate (dFdCTP), enabling incorporation into DNA and RNA during replication and inducing masked chain termination. Gem also inhibits ribonucleotide reductase (RR), thereby depleting intracellular deoxynucleotide pools and further impairing DNA synthesis (39). Gem has been reported to overcome oxaliplatin resistance and has been explored as an alternative strategy in oxaliplatin resistant CRC cases (40). Gem has also demonstrated superior efficacy over 5-FU in advanced pancreatic cancer (41).

In this project, it was hypothesized that modification of miR-15a with Gem would synergistically combine the tumor-suppressive effects of miR-15a with the chemotherapeutic activity of Gem, wherein all the cytidine molecules in the guide strand of miR-15a were substituted with Gem to develop the Gem-miR-15a mimic. Notably, Gem has been reported to exert immunomodulatory effects at low doses, including depletion of regulatory T cells in PDAC and CRC (42, 43). When released upon processing of Gem-miR-15a, by 5’ - 3’ exonucleases, released Gem may further modulate the tumor immune microenvironment (44). This multimodal design integrating multi-target gene regulation with direct cytotoxicity represents a unique therapeutic strategy. Importantly, Gem-miR-15a exhibits enhanced stability and can be delivered without a carrier vehicle, addressing key limitations inherent to conventional miRNA therapeutics.

In this study, the therapeutic potential of Gem-miR-15a was investigated using 5-FU- and oxaliplatin-resistant CRC cell lines and patient derived organoids, examining its effects on cell proliferation, apoptosis and cell cycle regulation. Gem-miR-15a demonstrated nanomolar efficacy in the resistant CRC models, effectively inducing cell cycle arrest and apoptosis, while suppressing multiple oncogenic drivers and pathways to overcome resistance. Most importantly, all effects were achieved under vehicle-free conditions. Collectively, this study provides mechanistic insight into the multimodal functionality of Gem-miR-15a, demonstrating its anti-tumor efficacy and underscoring its potential for clinical translation.

## Methods

### Synthesis of Gem-miR-15a mimics

Gem-miR-15a mimics were designed by substituting all the cytidine residues in the guide strand of miR-15a, with the nucleoside analog, Gem. No modifications were made to the passenger strand, to preserve target specificity and avoid potential off-target effects. The modified guide strand was then ordered separately along with the corresponding unmodified passenger strand from Dharmacon (Horizon Discovery). The oligonucleotides were purified using high-performance liquid chromatography (HPLC). The strands were then annealed at 60°C, prior to use.

### Cell lines and tissue culture

Human colorectal cancer cell lines (HCT116, HT-29, SW480 and SW620) were purchased from American Type Culture Collection (ATCC, Manassas, VA, USA). HCT116 and HT-29 cells were maintained in McCoy’s 5A (Modified) Medium (Thermo Fisher Scientific, Waltham, MA, USA) supplemented with 10% fetal bovine serum (FBS) (MilliporeSigma, St. Louis, MO, USA). SW480 and SW620 cells were maintained in Dulbecco’s Modified Eagle Medium (DMEM) (Thermo Fisher Scientific, Waltham, MA, USA) supplemented with 10% FBS. Prior to addition, FBS was sterilized via Steriflip-GP, 0.22 µm (MilliporeSigma, St. Louis, MO, USA).

The drug (5-FU, oxaliplatin)-resistant HCT116 cell lines were generated by repeated exposure of HCT116 cells to gradually increasing doses of 5-FU or oxaliplatin (MilliporeSigma, St. Louis, MO, USA). The cells were initially subjected to the IC_25_ of the drug for 48 hours in the respective growth media followed by replacement with drug-free media to allow recovery. This cycle was repeated with increasing concentrations of the drug, over several months until stable resistant cells were established: 5-FU Res HCT116 and Ox Res HCT116.

Spheroid forming SW620 cells were generated by plating SW620 cells in ultra-low attachment plates (Corning Costar 6 well plate) at 20,000 cells per well and were maintained in KnockOut™ DMEM/F12 medium supplemented with 20% KnockOut Serum Replacement (Thermo Fisher Scientific, Waltham, MA, USA), 1% L-glutamine (Thermo Fisher Scientific, Waltham, MA, USA), 10 ng/mL recombinant human basic fibroblast growth factor (bFGF), and 20 ng/mL recombinant human epidermal growth factor (rhEGF) (Biotechne, R&D systems, MN, USA). Growth factors were freshly added prior to media use. Cells were cultured in a humidified 5% CO□ incubator at 37°C for 5–7 days. Spheroid formation and morphology were monitored using a fluorescent microscope under bright-field conditions (10X and 20X objectives).

The p53 null, HCT116 (p53^-/-^, p21^+/+^) and p21 null, HCT116 (p53^+/+^, p21^-/-^) cells were obtained from the Bert Vogelstein Laboratory in Johns Hopkins School of Medicine. Both the cell lines were maintained in McCoy’s 5A (Modified) Medium (Thermo Fisher Scientific, Waltham, MA, USA) supplemented with 10% FBS (MilliporeSigma, St. Louis, MO, USA).

The luciferase-expressing colorectal cancer cell line, HCT116-Luc, was generated by transfecting the HCT116 with a luciferase-expressing lentiviral construct (obtained from Y. Ma laboratory at Renaissance School of Medicine at Stony Brook University). Post successful transfection, the cell line was maintained in McCoy’s 5A (Modified) Medium (Thermo Fisher Scientific, Waltham, MA, USA) supplemented with 10% fetal bovine serum (FBS) (MilliporeSigma, St. Louis, MO, USA). Luciferase expression was confirmed via *In Vivo* Imaging System (IVIS) before use in *in vivo* experiments.

### Transfection

For vehicle-mediated transfection, cells were seeded onto 6-well plates at a cell-density of 100,000 cells/well. 24 hours post-plating, cells were transfected with Oligofectamine (Thermo Fisher Scientific, Waltham, MA, USA) and the respective oligonucleotides, according to the manufacturer’s protocol, at a concentration of 50nM (negative control, miR-15a and Gem-miR-15a). The controls, pre-miR Negative Control #2 (non-specific scramble miRNA) and pre-hsa-miR-15a (unmodified miR-15a) were purchased from Thermo Fisher Scientific. Post 24-hours of transfection, Gem was added to the respective well at a concentration equivalent to the concentration of Gem in Gem-miR-15a (150nM). 5-hour post-transfection with oligonucleotides, media was changed to fresh media supplemented with 10% dialyzed fetal bovine serum (DFBS) (Thermo Fisher Scientific, Waltham, MA, USA).

For vehicle-free transfections, cells were seeded onto 6-well plates at a cell-density of 100,000 cells/well and 96-well plates at a density of 1000 cells/well. 24-hour post plating, media was changed with the corresponding oligonucleotide or drug dilutions, made in fresh media. The media was again changed with fresh media supplemented with 10% DFBS, 5 hours post transfection in 6-well plate and after 24 hours for 96-well plate.

### Cell proliferation assay

To investigate the cytotoxicity of Gem-miR-15a in the absence of a vehicle, cells were seeded onto a 96-well plate at 1000 cells/well in triplicates per concentration, followed by transfection after 24 hours, as described above. Cell viability was measured 6 days post-transfection using WST-1 Cell Proliferation Reagent (MilliporeSigma, St. Louis, MO, USA) or CellTitreGlo (Promega, Madison, WI, USA) as per manufacturer’s protocol on a SpectraMax iD3 plate reader (Molecular Devices, San Jose, CA, USA). For WST-1, cells were incubated in 10 μL WST-1 per 100 μL media for an hour at 37°C. Absorbance was then measured at 450 and 630 nm on SpectraMax. The O.D. was calculated by subtracting the absorbance at 630nm from that at 450 nm and the relative proliferation was calculated, after normalizing the O. . to untreated cells. For CellTitreGlo, 50 μL of CellTitreGlo was mixed with 50 μL of media and added to cells 6 days post-transfection. The contents were mixed for 2 minutes in an orbital shaker to induce cell lysis. The plate was then incubated at room temperature for 10 minutes and then luminescence was recorded on the SpectraMax iD3.

The IC_50_ values were calculated using GraphPad Prism 9 (GraphPad Software, San Diego, CA, USA), using the following equation:

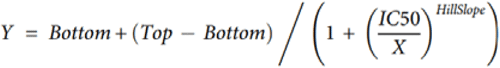

For the Spheroid cells (spSW620), the spheroids were dissociated into a single cell suspension and plated at 400 cells/well in 50 μL 10% Matrigel/hCGM in a 96 well plate, black, clear bottom (Falcon, Corning, NY, USA). 72-hours post-plating and visual confirmation of spheroid formation, they were treated with the desired concentrations of either Gem-miR-15a and 5-FU. After 6 days of treatment, cell viability was examined using CellTitre Glo, 3D (Promega, Madison, WI, USA) as per manufacturer’s protocol on a SpectraMax i3 plate reader (Molecular Devices, San Jose, CA, USA). 110 μL of CellTitreGlo 3D (equivalent to the volume of media in the well) was added to wells, followed by mixing the contents for 2 minutes in an orbital shaker to induce cell lysis. The plate was then incubated at room temperature for 30 minutes and then luminescence was recorded on the plate reader. The IC_50_ values were calculated using GraphPad Prism 9 (GraphPad Software, San Diego, CA, USA) as described above.

### Organoid culture and treatment

Patient-derived organoids HCRC4 (primary tumor, ascending colon), HCM-CSHL-0670-C18 (CRC670) (liver metastases, ascending colon) and HCM-CSHL-0726-C18 (CRC726) (liver metastases, upper rectum) were derived from the Stony Brook Medicine Biobank (Stony Brook Medicine Biobank, Renaissance School of Medicine, Stony Brook, USA). The organoid cells were cultured in Corning Matrigel GFR Membrane Matrix (Corning Incorporated, Corning, NY, USA) and incubated at 37°C in Human Complete Growth Medium (hCGM) containing the following: Advanced DMEM/F12 (Thermo Fischer Scientific, Waltham, MA, USA), HEPES (final concentration: 10 mM) (Thermo Fischer Scientific, Waltham, MA, USA), L-Glutamine (final concentration: 2 mM) (Thermo Fischer Scientific, Waltham, MA, USA), B-27 supplement (final concentration: 1X) (Thermo Fischer Scientific, Waltham, MA, USA), rhNoggin (final concentration: 100 ng/mL) (Biotechne, R&D systems, MN, USA), A 83-01 (final concentration, 500 nM) (Tocris Bioscience, Bristol, UK), SB202190 (final concentration, 10 μM) (Tocris Bioscience, Bristol, UK), rhEGF (final concentration: 50 ng/mL) (Biotechne, R&D systems, MN, USA), Gastrin I (human) (final concentration: 10 nM) (Tocris Bioscience, Bristol, UK), Nicotinamide (final concentration, 10 mM) (LKT Labs, MN, USA), N-acetylcysteine (final concentration, 1.25 mM) (LKT Labs, MN, USA).

For treatment with Gem-miR-15a and 5-FU, the organoids were dissociated into a single cell suspension and plated at 1000 cells/ well in 50 μL 10% Matrigel/hCGM in a 96 well plate, black, clear bottom (Falcon, Corning, NY, USA). 48-hours after plating and visual confirmation of organoid formation, they were treated with the desired concentrations of either Gem-miR-15a and 5-FU. After 6 days of treatment, cell viability was examined using CellTitreGlo, 3D (Promega, Madison, WI, USA) as per manufacturer’s protocol on a SpectraMax iD3 plate reader (Molecular Devices, San Jose, CA, USA). 110 μL of CellTitreGlo, 3D (equivalent to the volume of media in the well) was added to wells, followed by mixing the contents for 2 minutes in an orbital shaker to induce cell lysis. The plate was then incubated at room temperature for 30 minutes and the luminescence was recorded on the plate reader. The IC_50_ values were calculated using GraphPad Prism 9 (GraphPad Software, San Diego, CA, USA) as described above.

### Western immunoblot analysis

CRC cells were seeded and transfected with or without vehicle (oligofectamine), as described above. After 72 hours of transfection, cells were lysed using RIPA buffer (MilliporeSigma, St. Louis, MO, USA), mixed with a protease inhibitor cocktail (MilliporeSigma, St. Louis, MO, USA) and the proteins were collected for western immunoblot analysis. Proteins were probed with rabbit anti-WEE1 (Cell Signaling, 13084, 1:1,000), mouse anti-CHK1 (Cell Signaling, 2360, 1:1,000), rabbit anti-BMI1 (Cell Signaling, 6964, 1:1,000), rabbit anti-YAP1 (Cell Signaling, 4912, 1:1,000), rabbit anti-CCND1 (Cell Signaling, 55506, 1:1,000), rabbit anti-p53 (Cell Signaling, 9282, 1:1,000), mouse anti-p21 (Cell Signaling, 2946, 1:2,000), rabbit anti-CCNB1 (Cell Signaling, 12231, 1:1,000), rabbit anti-CCNA2 (Cell Signaling, 67955, 1:1,000), rabbit anti-Phospho-Histone H2A.X (γH2AX) (Cell Signaling, 2577, 1:1,000), mouse anti-thymidylate synthase (Sigma Aldrich, MAB4130, 1:1,000), rabbit anti-BAK (Cell Signaling, 12105, 1:1,000), rabbit anti-PARP (Cell Signaling, 9532, 1:1,000), rabbit anti-cleaved PARP (Cell Signaling, 5625, 1:1,000), rabbit anti-caspase-7 (Cell Signaling, 12827, 1:1,000), rabbit anti-cleaved caspase-7 (Cell Signaling, 8438, 1:1,000) and mouse anti-β-actin (Merck, A5441, 1:10,000). After probing with primary antibodies, which were diluted in 5% milk (Bio-Rad, Hercules, CA, USA) made in TBST, the proteins were incubated with secondary antibodies, either goat anti-rabbit-HRP (Bio-Rad, 1721019, 1:5,000) or goat anti-mouse-HRP (Bio-Rad, 1706516, 1:5,000), depending on the corresponding primary antibody. The protein bands were then visualized on the LI-COR Biosciences Odyssey Fc imaging system, after incubating with SuperSignal West Pico PLUS Chemiluminescent Substrate (Thermo Fisher Scientific, Waltham, MA, USA). The bands were then quantified and analysed using Image Studio Version 5.2.5 (LI-COR Biosciences, Lincoln, NE, USA).

### Cell cycle assay

CRC cells were seeded and transfected with the respective oligonucleotides along with vehicle (oligofectamine) as described above or treated with 150 nM Gem (equimolar to the Gem present in Gem-miR-15a). After 48 hours of transfection, the cells were resuspended in Krishan modified buffer, supplemented with 0.02 mg/mL RNase H (Thermo Fisher Scientific, Waltham, MA, USA) and 0.05 mg/mL propidium iodide (MilliporeSigma, St. Louis, MO, USA). Cells were analysed by flow cytometry in the CytoFLEX Flow Cytometer (Beckman Coulter, Brea, CA, USA) and the results were analysed using Modfit LT Software (BD Biosciences, Sparks, MD, USA) to determine the number of cells in G1, S and G2 phases of cell cycle.

### Apoptosis assay

CRC cells were seeded and transfected with vehicle (oligofectamine) as described above and after 72 hours of transfection, cells were collected and stained with Annexin V (Thermo Fisher Scientific, Waltham, MA, USA) and propidium iodide (MilliporeSigma, St. Louis, MO, USA). The stained cells were then analysed by flow cytometry in the CytoFLEX Flow Cytometer (Beckman Coulter, Brea, CA, USA) and the Annexin V-positive cells were categorized as apoptotic.

### RNA sequencing

CRC cells were seeded and transfected without vehicle as described above and 72-hours post transfection, RNA was extracted using TRIzol reagent (Thermo Fisher Scientific, Waltham, MA, USA), according to the manufacturer’s protocol. RNA was isolated from three different biological replicates per condition and pooled prior to sequencing. The RNA samples were subsequently submitted to Novogene (Sacramento, CA, USA) for Human mRNA Sequencing (WBI-Quantification). The sequencing data was then provided to the Ramana lab (Department of Biomedical Informatics, Stony Brook University) for bioinformatic analysis and generation of heatmaps. The pathway enrichment analysis and resistance reversal analysis were performed using R-programming (v4.2). In R, gene set enrichment analysis (GSEA) was conducted using the clusterProfiler package, with genes ranked based on log2 fold-change values for each comparison (Par and 5-FU Res conditions). GSEA was performed using KEGG pathway gene sets with the gseKEGG function, based on the original GSEA framework. Functional enrichment analyses were conducted using KEGG pathways and Gene Ontology (GO) annotations.

A pathway “reversion” score was defined as:

Reversion score = −(NES_resistance × NES_G15a in 5-FU Res HCT116) where,

NES_resistance represents the normalized enrichment score obtained from the comparison of untreated 5-FU-resistant cells versus untreated parental cells (5-FU Res NC vs Par NC), reflecting resistance-associated pathway enrichment.

All analyses and visualizations were performed using R, and scripts are available on request.

### Metastatic CRC mouse model

All animal procedures were performed upon approval from the Stony Brook University Institutional Animal Care and Use Committee (IACUC). Non-obese diabetic (NOD)/ severe combined immunodeficiency (SCID) mice (JAX: 001303) were purchased from The Jackson Laboratory (The Jackson Laboratory, Bar Harbour, ME, USA). Eight-week-old NOD/SCID mice (2 males, 2 females per treatment group) were injected with 2 x 10^6^ HCT116 (+Luc) cells suspended in 0.2 mL PBS, intravenously (IV) via the tail-vein. After confirmation of successful engraftment (2 weeks from injection), mice were randomized into three groups (Vehicle control, Gem-miR-15a and Gem), where they were treated with either 80µg of PEI-MAX (vehicle control) (Polysciences, Warrington, PA, USA) in 5% D-glucose or 80µg of PEI-MAX along with 80µg of Gem-miR-15a (4 mg/kg) in 5% D-glucose (Gem-miR-15a), or 50 mg/kg of Gem in 1X DPBS. The Vehicle control and Gem-miR-15a groups were treated on alternate days for 2 weeks (eight times) via IV injections. The Gem group was treated every three days for 2 weeks (four doses) via intraperitoneal (IP) injections. To monitor tumor advancement, mice were injected with IVIS Brite RediJect D-Luciferin (PerkinElmer) and after 10 minutes luciferase expression was measured using the IVIS spectrum In Vivo Imaging System (IVIS) (PerkinElmer). Tumor growth was measured as a function of (total flux at time of measurement [p/s])/ (total flux at initial measurement [p/s]). Weights of mice were monitored daily and euthanized according to IACUC protocol (>15 % weight loss over 7 days).

### Statistical analysis

All experiments were conducted at least three times and analysed using GraphPad Prism 9 (GraphPad Software, San Diego, CA, USA). Statistical significance between 2 groups was determined using the student’s t test. For the *in vivo* studies, two-way ANOVA with mixed effects was performed to evaluate the impact of treatment over time on the tumor progression rate in mice. Data was presented as mean ± SD. A p-value of <0.05 was considered to be statistically significant (*p<0.05, **p<0.01, ***p<0.001, ****p<0.0001).

ChatGPT (Open AI) was used solely to improve language quality and clarity during manuscript preparation. The authors reviewed, edited and approved all AI-assisted text and take full responsibility for the final content of the manuscript.

## Results

### Gem-miR-15a exhibits enhanced anti-proliferative efficacy in CRC *in vitro*, under vehicle-free conditions

The efficacy of Gem-miR-15a was investigated across various CRC cell lines, including HCT116 (p53^+/+^, p21^+/+^), HT-29 (p53^R273H/mut^, p21^+/+^), SW480 (p53^mut^, p21^+/+^), SW620 (p53^mut^, p21^+/+^, metastatic derivative of SW480), as well as HCT116 (p53^-/-^, p21^+/+^) and HCT116 (p53^+/+^, p21^-/-^) cells. As mentioned earlier, Gem-miR-15a was generated by substituting all the cytidine residues on the guide strand of miR-15a with Gem, while the passenger strand was left unmodified (Figure 1A).

**Figure 1:**
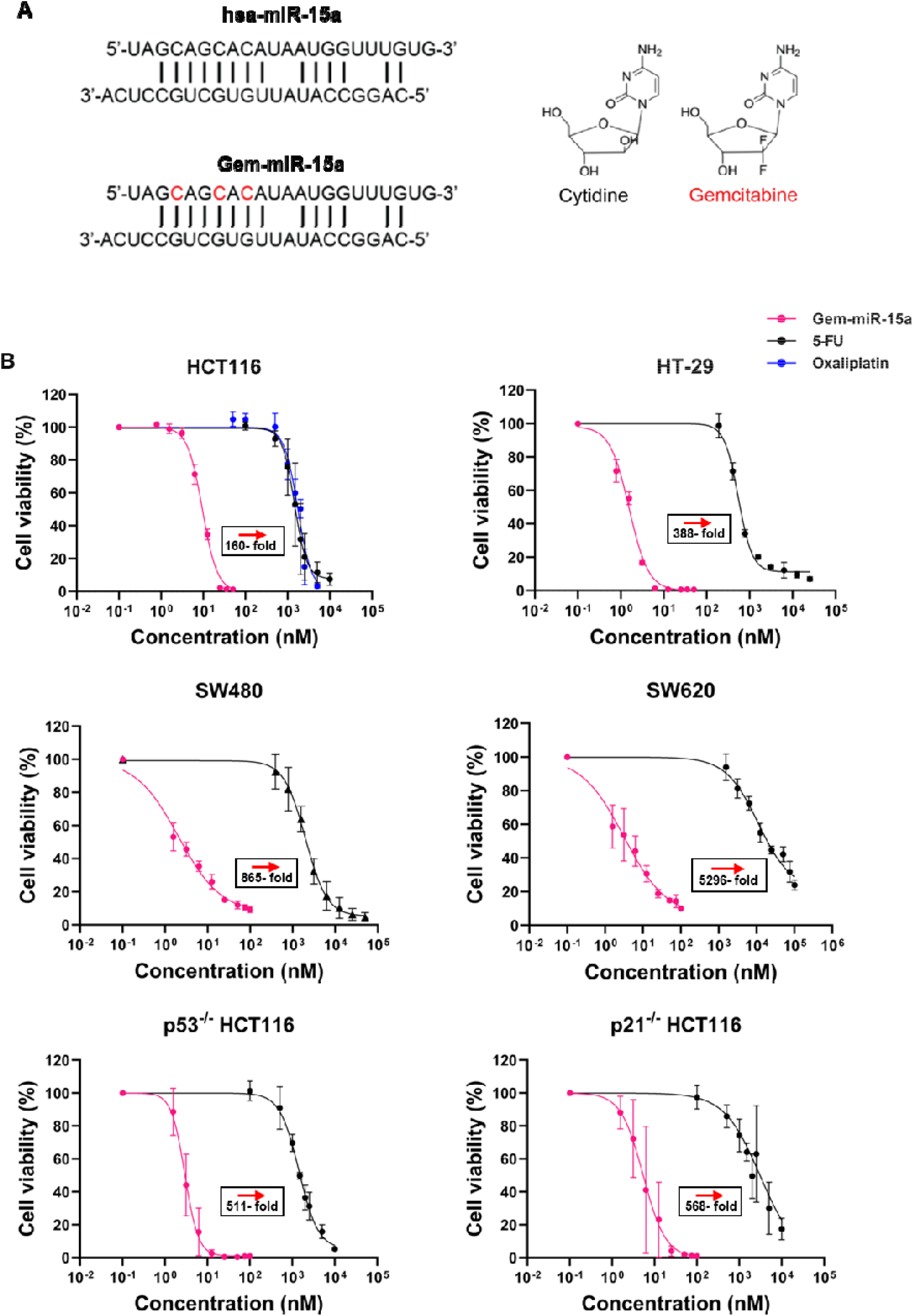
Gem-miR-15a exhibits potent anti-proliferative efficacy across multiple colorectal cancer (CRC) cell lines. (A) Schematic representation of Gem-miR-15a design, in which cytidine residues in the guide strand of hsa-miR-15a are substituted with nucleoside analog gemcitabine (red), while the passenger strand remains unmodified. (B) Dose-response analysis demonstrating the effects of Gem-miR-15a on cell viability across multiple CRC cell lines, including HCT116, HT29, SW480, SW620 as well as p53^-/-^ and p21^-/-^ HCT116 cells (n=3). Gem-miR-15a (pink) shows markedly enhanced anti-proliferative compared to 5-fluorouracil (5-FU, black) and oxaliplatin (blue). Cells were treated with increasing concentration of each drug and viability was assessed after 6 days using WST-1 assay. Gem-miR-15a was delivered without the use of a transfection vehicle. IC_50_ values are listed in Table 1. Data are represented as mean ± SD from at least three independent experiments.

A dose-dependent reduction in cell viability was observed following treatment with Gem-miR-15a across all the cell lines, with half maximal inhibitory concentration (IC_50_) values in the nanomolar range (1.53 – 9.47 nM) (Figure 1B, Table 1). In contrast, 5-fluorouracil (5-FU) and oxaliplatin exhibited much higher IC_50_ values (micromolar range), indicating substantially lower potency as compared to Gem-miR-15a (∼160 – 5000-fold enhancement) (Table 1). Similar nanomolar efficacy was observed in the HCT116 (p53^-/-^, p21^+/+^) and HCT116 (p53^+/+^, p21^-/-^) cells, suggesting that the antiproliferative effects of Gem-miR-15a are independent of the p53 pathway. Notably, Gem-miR-15a was delivered in the absence of a delivery vehicle, demonstrating intrinsic cellular uptake. A dose response assay was also performed for the native counterpart, miR-15a in HCT116 cells, which showed an IC_50_ value of 130.46 nM (Supplementary Figure 1). A delivery vehicle was used for the transfection of miR-15a, since it cannot cross the lipid bilayer of the cell membranes, hindering its cellular uptake.

**Table 1:**
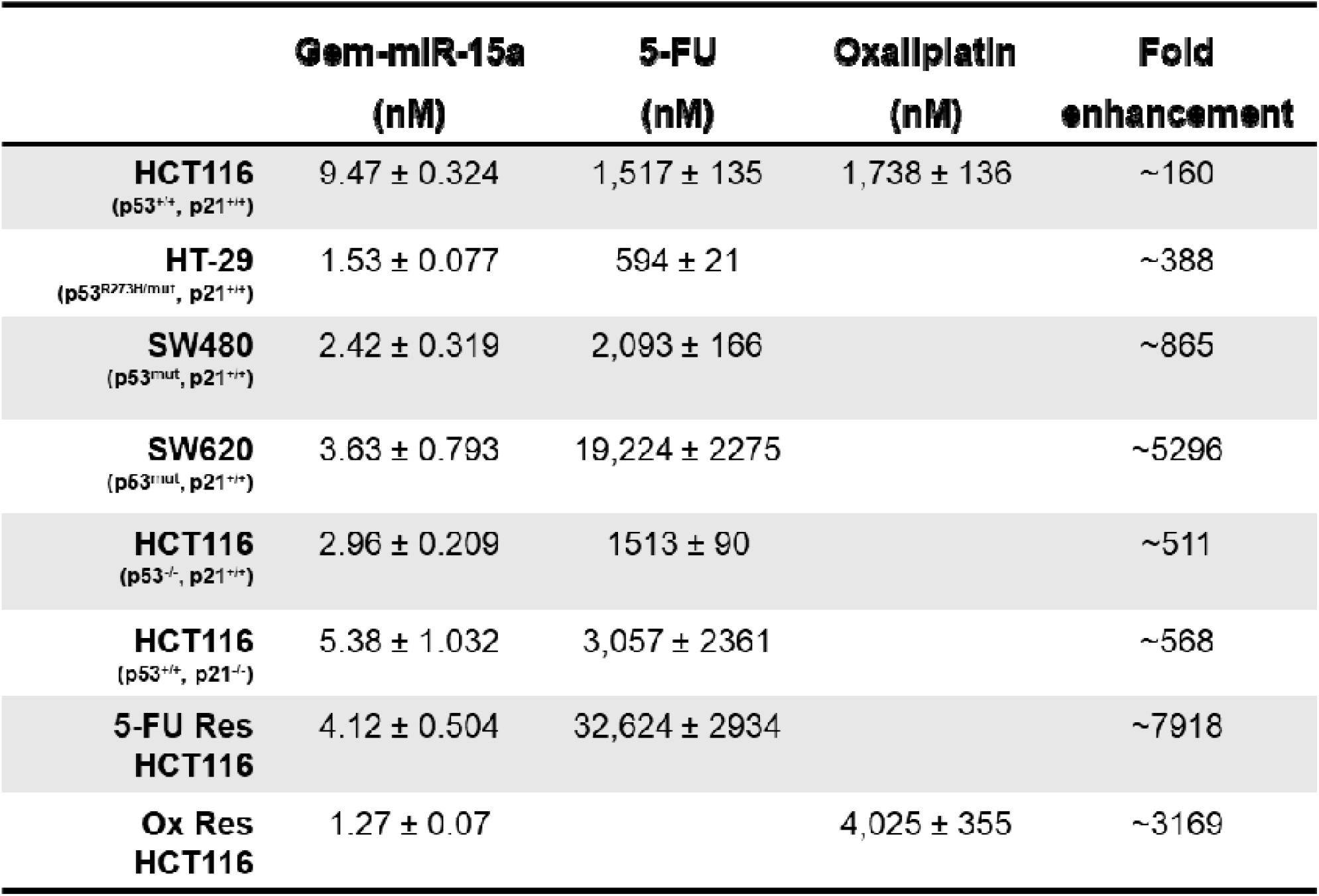
IC_50_ values of Gem-miR-15a and 5-FU in CRC cell lines. Half-maximal inhibitory concentrations (IC_50_) values of Gem-miR-15a, 5-fluorouracil (5-FU) and oxaliplatin (in HCT116 cells) across multiple CRC cell lines. IC_50_ values were calculated using non-linear regression analysis in GraphPad Prism. Data are represented as mean ± SD, from at least three independent experiments.

### Gem-miR-15a effectively induces S-phase cell cycle arrest, abolishes cells in G2 and induces apoptosis across CRC cells, under vehicle-free conditions

To further investigate the mechanism underlying growth inhibition, cell cycle distribution was analyzed by propidium iodide (PI) staining. A pronounced loss of G2-phase cells and accumulation in S-phase was observed following Gem-miR-15a treatment across all CRC cell lines, as reflected by a significant reduction in G2/S ratio as compared to the negative control (Figure 2A). The effects were markedly greater than that observed in the control conditions, including unmodified miR-15a or Gem alone. For instance, in HCT116 and HT-29 cells, Gem-miR-15a treatment resulted in a significant reduction in G2/S ratio from 1 in control cells, to near zero, following treatment (p<0.0001). A similar pattern of enhanced cell cycle disruption was consistently observed across additional CRC cell lines, including SW480 (p=0.0006), SW620 (p<0.0001), as well as in HCT116 (p53^-/-^, p21^+/+^) (p<0.0001) and HCT116 (p53^+/+^, p21^-/-^) (p=0.0009) cells.

**Figure 2:**
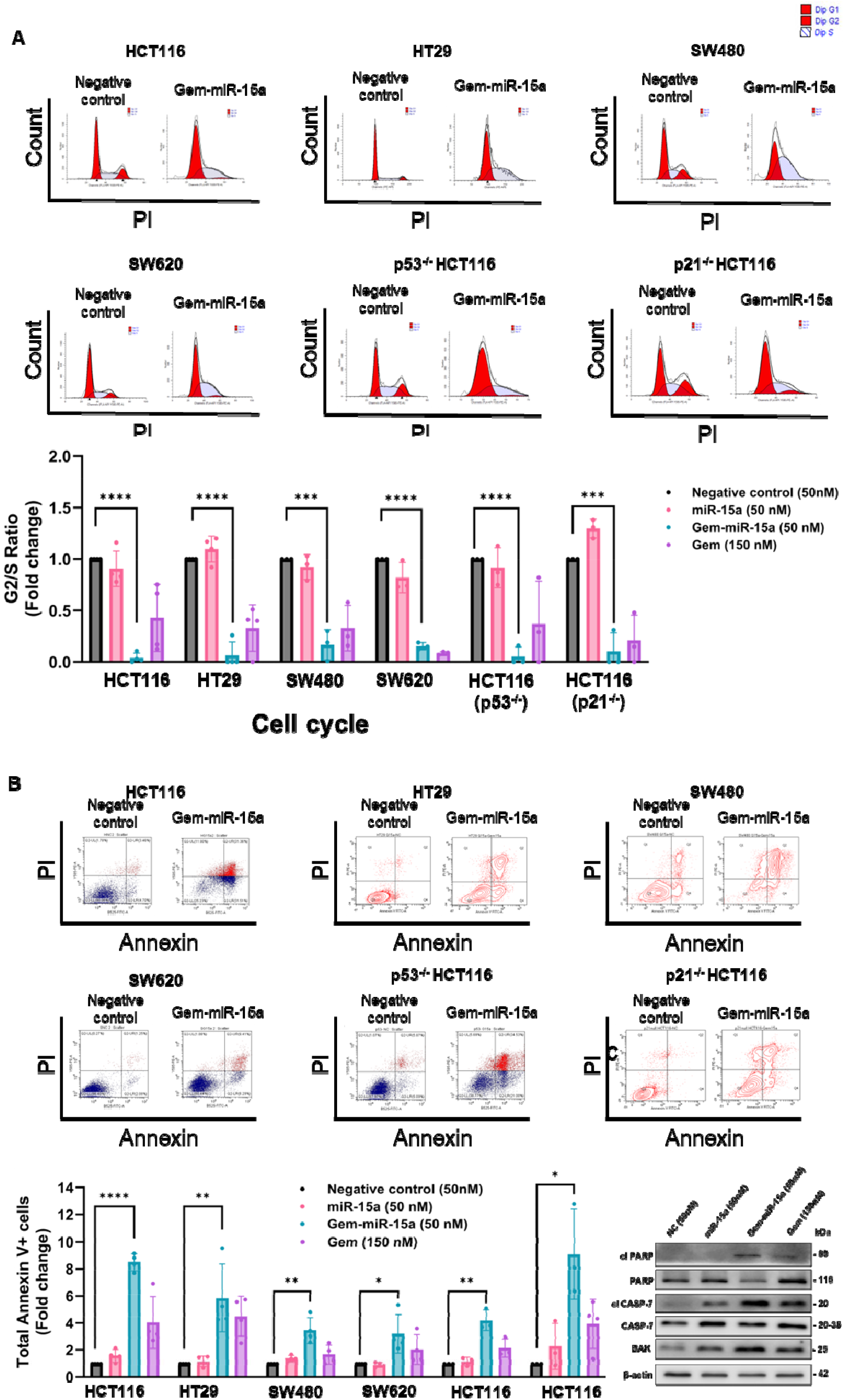
Gem-miR-15a induces cell cycle arrest and apoptosis in CRC cells. (A) Cell cycle analysis following Gem-miR-15a treatment across CRC cell lines (HCT116, HT29, SW480, SW620, p53^-/-^ and p21^-/-^ HCT116 cells; n=3). Representative flow cytometry profiles show a marked reduction in the G2-cell population. Quantification of G2/S ratio demonstrates a significant decrease in Gem-miR-15a treated cells, compared to negative control (NC), consistent with S-phase arrest. (B) Apoptosis analysis by Annexin V/PI staining showing increased apoptotic cells following Gem-miR-15a treatment. Quantification of total Annexin V-positive cells (fold change, n=3) demonstrates a significant increase compared to NC. (C) Western blot analysis confirming apoptosis induction, as indicated by increased expression of cleaved caspase-7, cleaved PARP and BAK. Cells were treated with 50 nM of NC, miR-15a, Gem-miR-15a or 150 nM of gemcitabine (Gem). Data are represented as mean ± SD from at least three independent experiments. Statistical comparisons were performed between NC and Gem-miR-15a groups using Student’s t-test. *p<0.05, **p<0.01, ***p<0.001, ****p<0.0001.

Additionally, the effects on apoptosis were evaluated using Annexin V/PI staining. A significant increase in Annexin V-positive cells was observed following Gem-miR-15a treatment across all cell lines with an ∼8-fold increase in HCT116 cells (p ≤ 0.0001) and a ∼4-6-fold increase in HT-29, SW480 and SW620 (p =0.0084, p =0.0022 and p =0.023 respectively). The apoptosis induction was most prevalent in Gem-miR-15a treated cells as compared to all other conditions. The results were similar for HCT116 (p53^-/-^, p21^+/+^) and HCT116 (p53^+/+^, p21^-/-^) cells, with an apoptotic increase by 4-fold and 8-fold, as compared to the negative control (p = 0.002 and p = 0.01 respectively). The representative Annexin V/propidium iodide flow cytometry plots also show an increase in apoptotic cells in the Gem-miR-15a treated cells as compared to the respective controls (Figure 2B, Supplementary Figure 3). These findings were supported by western blot analysis, which demonstrated increased expression of cleaved PARP, cleaved caspase-7 and BAK.

Collectively, these results indicate that that Gem-miR-15a suppresses proliferation through coordinated induction of S-phase arrest and apoptosis in CRC cells.

### Gem-miR-15a overcomes chemoresistance (5-FU, Oxaliplatin) in resistant CRC cells

Resistance to 5-FU-based chemotherapies, such as FOLFOX is the main cause of treatment failure and recurrence in CRC (45).

To investigate the effect of Gem-miR-15a on acquired chemoresistance, 5-FU resistant HCT116 (5-FU Res HCT116) and Oxaliplatin resistant HCT116 (Ox Res HCT116) were generated by repeated exposure of HCT116 cells to increasing dose of 5-FU and Ox for a continued time, progressively selecting for the cells that grew in the presence of the drugs, that were then used for further testing. A robust dose-dependent reduction in cell viability was observed across the resistant CRC models, with IC_50_ values in the low nanomolar range (4.12 nM in 5-FU Res HCT116 and 1.27 nM in Ox Res HCT116), whereas the corresponding standard therapies exhibited markedly reduced efficacy. Compared to the corresponding standard therapy, Gem-miR-15a showed a ∼3000 and 8000-fold enhancement in IC_50_ (Figure 3A, Table 1).

**Figure 3.**
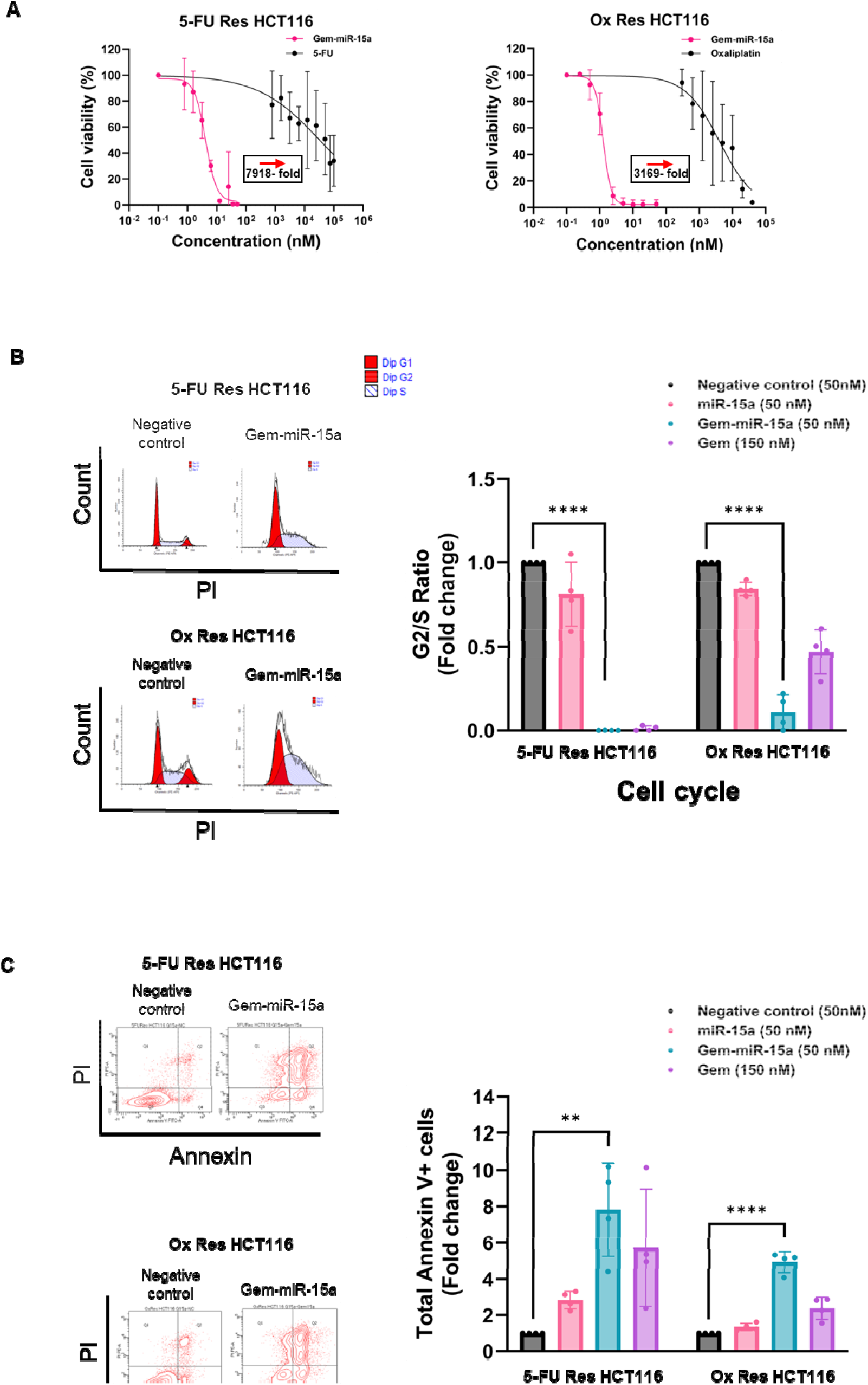
Gem-miR-15a overcomes chemoresistance and retains efficacy in resistant CRC models. (A) Dose-response analysis demonstrating the effects of Gem-miR-15a on cell viability across resistant CRC models, including 5-FU-resistant (5-FU Res) and oxaliplatin-resistant (Ox Res) HCT116 cells (n=3). Gem-miR-15a (pink) exhibits markedly enhanced anti-proliferative activity compared to corresponding treatments: 5-FU (black) in 5-FU Res HCT116 cells and oxaliplatin (black) in Ox Res HCT116 cells. IC_50_ values are listed in Table 1. (B) Cell cycle analysis of resistant CRC models following Gem-miR-15a treatment. Representative propidium (PI) staining profiles show a reduction in the G2-cell population. Quantification of G2/S ratio (fold change, n=3) demonstrates a significant decrease in Gem-miR-15a treated cells compared to NC, consistent with S-phase arrest across resistant models. (C) Apoptosis analysis by Annexin V/PI staining showing increased apoptotic cells following Gem-miR-15a treatment. Representative plots and quantification of total Annexin V-positive cells (fold change, n=3) demonstrate a significant increase in apoptosis compared to NC across all three resistant cell types. Cells were treated with 50 nM of NC, miR-15a, Gem-miR-15a or 150 nM of gemcitabine (Gem). Data are represented as mean ± SD from at least three independent experiments. Statistical comparisons were performed between NC and Gem-miR-15a groups using Student’s t-test. *p<0.05, **p<0.01, ***p<0.001, ****p<0.0001.

Consistent with parental cells, a significant reduction in the G2/S ratio was observed following treatment compared to the negative control in the resistant models (p ≤ 0.0001 for 5-FU Res HCT116 and Ox-Res HCT116), with an increased number of cells in S phase and almost no cells in G2 for both the resistant cell lines, indicating S-phase cell cycle arrest (Figure 3B). In parallel, apoptosis analysis revealed a marked increase in the Annexin V-positive population following treatment (p = 0.0018 for 5-FU Res HCT116 and p ≤ 0.0001 for Ox-Res HCT116) (Figure 3C). Notably, Gem-miR-15a synergized with 5-FU in 5-FU Res HCT116 cells, reflected by a HSA synergy score of 11.78 and CSS of 66.36, further supporting its potential to restore sensitivity to fluoropyrimidine-based therapy (Supplementary Figure 4).

Consequently, these findings further support that Gem-miR-15a can effectively overcome resistance to both the standard chemotherapies.

### Gem-miR-15a retains target specificity and downregulates key oncogenic targets in CRC cells

To ensure Gem-miR-15a maintains target specificity post-modification with Gem, and suppresses key oncogenic targets, protein expression of some miR-15a targets was evaluated by western blot analysis in HCT116 and HT-29 cells. A reduction in the expression of established miR-15a targets, including WEE1 (p=0.001 in HCT116, p=0.01 in HT-29), CHK1 (p=0.0004 in HCT116), YAP1 (p=0.01 in HCT116, p=0.003 in HT-29), BMI1 (p=0.006 in HCT116, p=0.0008 in HT-29) and CCND1 (p=0.0002 in HT-29) was observed following Gem-miR-15a treatment (Figure 4A, B), confirming retention of target specificity. The downregulation was even more pronounced in the Gem-miR-15a treated cells as compared to miR-15a treated cells. Similar target suppression was observed in the 5-FU Res HCT116 cells (Supplementary Figure 2).

**Figure 4.**
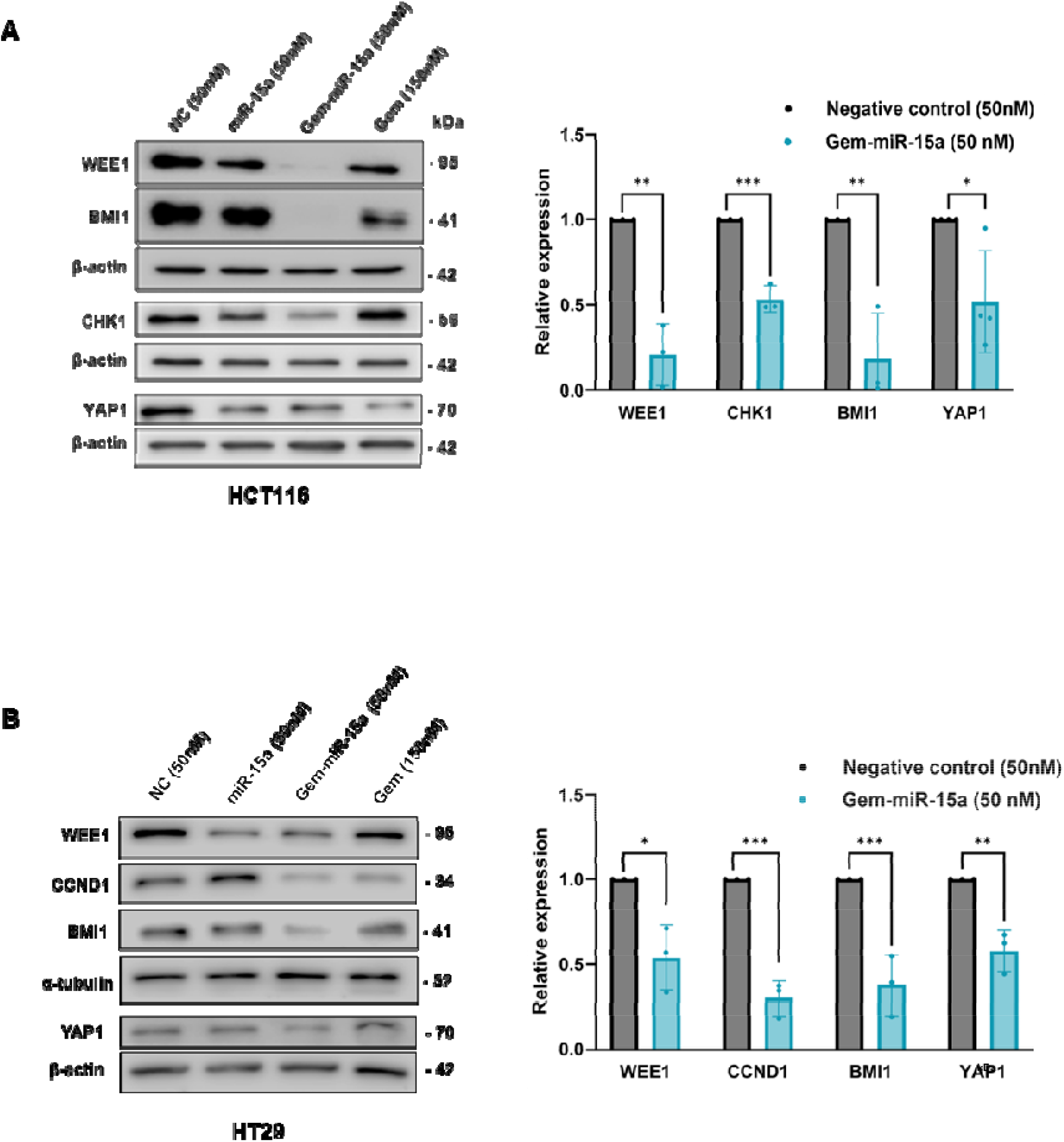
Gem-miR-15a retains target specificity and suppresses oncogenic proteins. (A, B) Western blot analysis of HCT116 and HT29 cells treated with NC, miR-15a, Gem-miR-15a or gemcitabine (Gem), demonstrating expression of established miR-15a target proteins. Key oncogenic targets, including WEE1, CHK1, BMI1, YAP1 and CCND1, are shown. Gem-miR-15a treatment results in consistent downregulation of these targets, compared to NC, indicating retention of target specificity following chemical modification. Corresponding densitometric quantification comparing NC and Gem-miR-15a-treated groups is presented. Data are represented as mean ± SD from at least three independent experiments. Statistical comparisons were performed between NC and Gem-miR-15a groups using Student’s t-test. *p<0.05, **p<0.01, ***p<0.001, ****p<0.0001.

### Gem-miR-15a modulates gene expression and signaling pathways in CRC, supported by RNA-seq and protein analysis

To elucidate the molecular mechanism by which Gem-miR-15a works and overcomes resistance to chemotherapy, RNA-seq analysis was conducted on parental HCT116 and 5-FU Res HCT116 cells, each under untreated and Gem-miR-15a treated conditions. KEGG pathway enrichment analysis revealed modulation of pathways associated with apoptosis, inflammatory signaling, p53, mTOR, autophagy and transcriptional regulation in both parental and 5-FU Res HCT116 (Figure 5A), many of which have been previously implicated in CRC progression and chemoresistance. Further, to quantify the reversal of resistance-associated pathways, a pathway reversal score was calculated. Gem-miR-15a treatment efficiently reversed multiple 5-FU resistance-associated pathways. It upregulated various inflammatory and stress-induced pathways such as the cytokine-cytokine receptor pathway, PI3K-Akt and NOD-like receptor pathway, while suppressing autophagy, HIF-1, Hippo and mTOR pathways, which are known to contribute to tumor survival, metabolic adaptation and resistance to fluoropyrimidine-based therapies. Additionally, at the gene level, the quadrant plot demonstrates bidirectional correction/ reversal of the resistance associated expression patterns by Gem-miR-15a, with downregulation of genes such as *AKT3, MAP2K6* and *WNT16* that were upregulated in resistant cells, consistent with prior studies linking PI3K-Akt and MAPK signaling to chemoresistant CRC, while upregulation of genes such as *MUC2* and *CASP10* that were suppressed in the resistant cells, suggesting restoration of epithelial differentiation and apoptotic signaling pathway that are often compromised in resistant tumors (Figure 5B).

**Figure 5.**
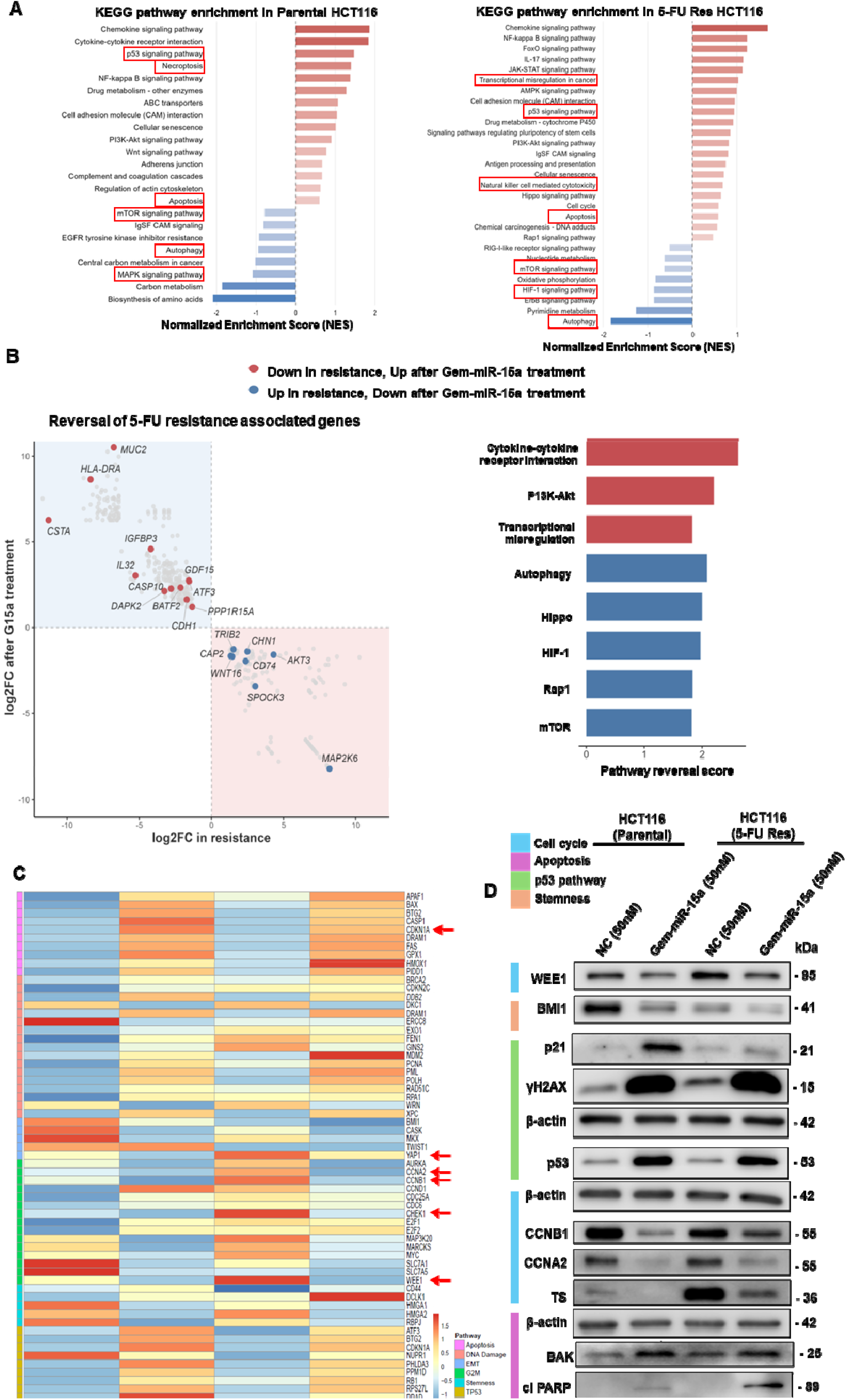
Gem-miR-15a modulates signaling pathways and reverses resistance-associated programs at the RNA and protein level. (A) KEGG pathway enrichment analysis of differentially expressed genes following Gem-miR-15a (G15a) treatment in parental and 5-FU Res HCT116 compared to NC. Pathways are shown based on normalized enrichment score (NES), with upregulated pathways in red and downregulated pathways in blue. (B) Resistance reversal analysis in 5-FU Res HCT116 following G15a treatment. Bar plot showing pathway reversal scores, highlighting pathways reversed by G15a. A corresponding quadrant plot illustrates gene-level expression changes, with genes upregulated in resistance and downregulated after treatment shown in blue, and genes downregulated in resistance and upregulated after treatment shown in red. (C) Heatmap of differentially expressed genes across parental and 5-FU Res HCT116 cells under untreated and G15a-treated conditions. Genes are grouped by functional pathways, demonstrating a shift in expression profiles of resistant cells toward a treated parental like state, after treatment. Selected genes validated at the protein level are highlighted. (D) Western blot analysis of parental and 5-FU Res HCT116 cells under untreated and G15a-treated conditions, showing modulation of proteins involved in cell cycle regulation, DNA damage response, stemness and apoptosis. G15a treatment results in downregulation of oncogenic targets and upregulation of p53 pathway and apoptosis markers. RNA was extracted from biological replicated (n=3) and pooled for sequencing. Western blot data are representative of at least three independent experiments.

Further, a heatmap analysis of differential gene expression revealed distinct gene expression profiles between untreated and Gem-miR-15a treated cells across both parental and resistant cells, with the treated resistant cells exhibiting similar expression profiles with the treated parental cells (Figure 5C, Supplementary figure 5). In the treated parental and 5-FU Res HCT116 cells, upregulation of key apoptotic genes including *APAF1, BAX and FAS* were observed. In parallel, genes associated with DNA damage response, including *RAD51C, DDB2, XPC* and *EXO1*, suggest activation of repair pathways following Gem-miR-15a treatment. Additionally, downregulation of *AURKA*, a key regulator of mitotic progression and cell proliferation, was observed, which has been previously associated with enhanced sensitivity to chemotherapeutic agents in CRC, supporting cell cycle disruption. Consistent with the activation of p53 pathway, key downstream targets including *CDKN1A* (p21), *BTG2* and *SESN1* were upregulated following treatment, further indicating activation of stress-response and cell cycle arrest programs (Figure 5C).

Western blot analysis further validated the RNA-seq results in both parental and resistant cells demonstrating a modulation of pathways associated with cell cycle regulation, DNA damage response and apoptosis. In addition to reduction in established targets, WEE1, BMI1 and YAP1 in both parental vs 5-FU Res HCT116 cells, there was also an increase in the expression of p53, p21 and γH2AX, indicating DNA damage and p53 pathway induction. Additionally, an increased expression of cleaved caspase-7, cleaved PARP and BAK, suggested an induction of apoptosis. Interestingly, Gem-miR-15a treatment reduced the expression of cyclins involved in cell cycle progression such as CCNB1 and CCNA2 in both parental and 5-FU Res HCT116 cells (Figure 5D). Notably, thymidylate synthase (TS), which was elevated in untreated 5-FU resistant cells and is a known mechanism of 5-FU resistance, was suppressed following Gem-miR-15a treatment. Several genes corresponding to the protein level changes including *WEE1, CHK1, CDKN1A* and *YAP1* are highlighted in the heatmap (Figure 5C), demonstrating concordance between transcriptomic and protein analyses.

Together, these results demonstrate that Gem-miR-15a not only inhibits resistance-associated pathways but also reverses critical regulatory networks, by modulating the expression of genes as well as proteins, thereby explaining its role in overcoming acquired resistance.

### Gem-miR-15a exhibits robust antitumor activity in clinically relevant 3D CRC models and synergizes with oxaliplatin

To evaluate the therapeutic efficacy of Gem-miR-15a in physiologically relevant models, its activity was assessed in 3D CRC models, such as spheroid forming SW620 cells, and multiple patient-derived organoids, both primary and metastatic (HCRC4, CRC670 and CRC726). A dose-dependent reduction in viability with IC_50_ values ranging from 6.17 – 13.43 nM, was observed in the 3D models, whereas 5-FU exhibited substantially lower potency (∼100-1000 fold decreased relative potency) (Figure 6A, Table 2). Consistent with these findings, morphological analysis under brightfield microscopy, revealed progressive loss of spheroid and organoid integrity with increasing concentrations of Gem-miR-15a (Figure 6B). Western blot analysis in organoid CRC670 confirmed the suppression of key targets WEE1 (p=0.004) and CHK1 (p=0.003) (Figure 6C). Lastly, to evaluate the combinatorial potential of Gem-miR-15a with oxaliplatin in HCT116 cells drug combination analysis was done, which revealed synergistic interaction between the two drugs with a mean Bliss synergy score of 13.99. The combination also exhibited a high combination sensitivity score (CSS) of 87.56, exceeding the individual response indices for Oxaliplatin (RI_1_ =16.02) and Gem-miR-15a (RI_2_ =39.13). The synergy heatmap illustrating the multiple dose combinations reveals the enhanced cytotoxic effects of the combination as compared to either drug alone. These findings indicate that Gem-miR-15a potentiates the therapeutic efficacy of oxaliplatin (Figure 6D).

**Figure 6.**
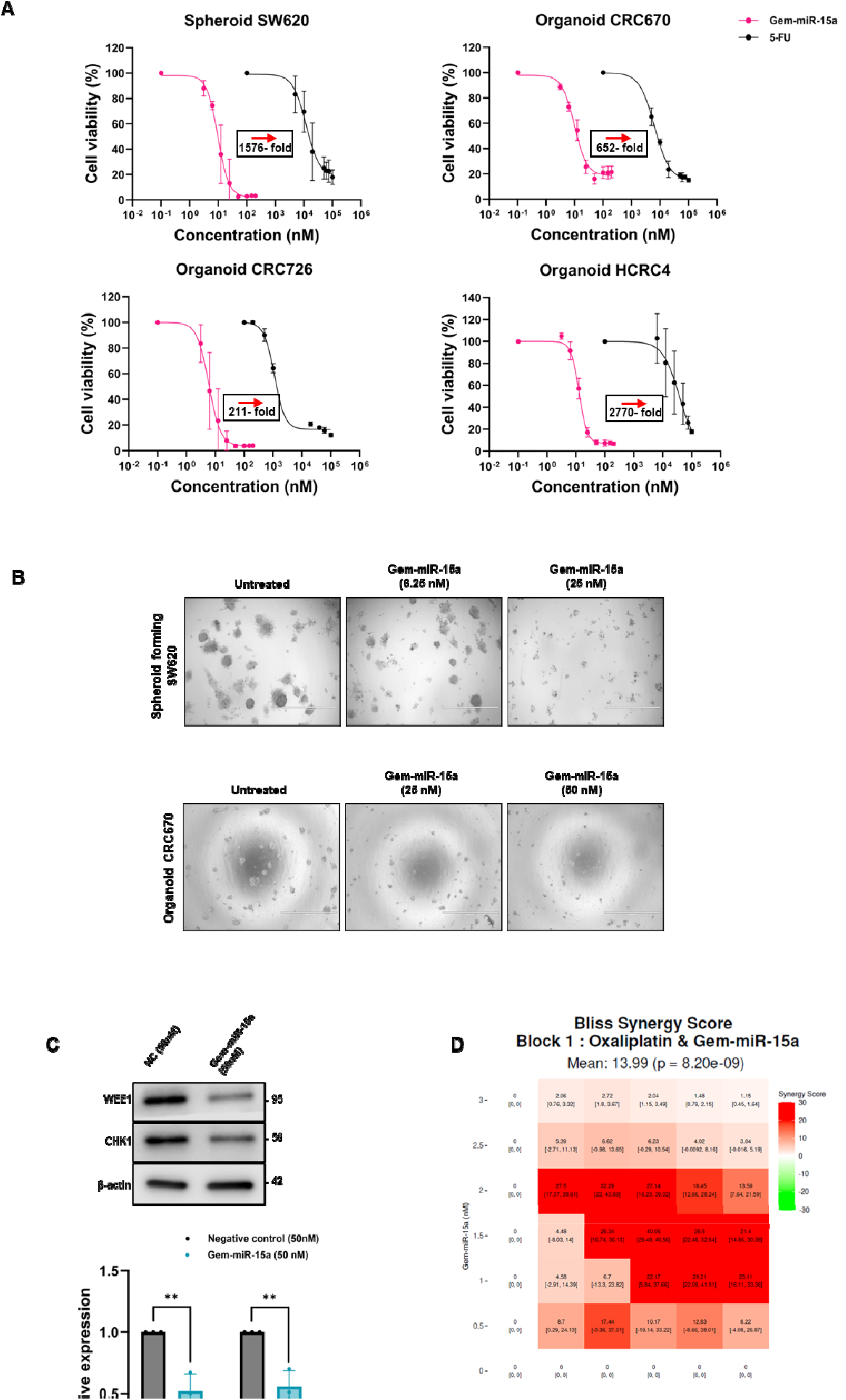
Gem-miR-15a demonstrates efficacy in 3D models and synergizes with oxaliplatin. (A) Dose-response analysis demonstrating the effect of Gem-miR-15a on cell viability in 3D CRC models including spheroid-forming SW620 cells and patient-derived organoids (HCRC4, CRC670, CRC726) (n=3). Gem-miR-15a (pink) exhibits enhanced anti-proliferative activity compared to 5-FU (black). IC_50_ values are listed in Table 2. (B) Representative images of SW620 spheroids and CRC670 organoids showing dose-dependent morphological disruption and loss of structural integrity following Gem-miR-15a treatment. (C) Western blot analysis of CRC670 organoids treated with NC or Gem-miR-15a, demonstrating downregulation of WEE1 and CHK1. Corresponding densitometric quantification comparing NC and Gem-miR-15a groups is shown. (D) Synergy analysis of Gem-miR-15a in combination with oxaliplatin in parental HCT116 cells. Heatmap representation of drug interactions shows a Bliss synergy score of 13.99, indicating synergistic effect. Data are derived from dose-response matrix analysis using SynergyFinder+. Data are represented as mean ± SD from at least three independent experiments. Statistical comparisons were performed between NC and Gem-miR-15a groups using Student’s t-test. *p<0.05, **p<0.01, ***p<0.001, ****p<0.0001.

**Table 2:** IC_50_ values of Gem-miR-15a and 5-FU in 3D CRC models. Half-maximal inhibitory concentrations (IC_50_) values of Gem-miR-15a and 5-fluorouracil (5-FU) in three-dimensional (3D) CRC models, including spheroid-forming SW620 and patient-derived organoids. IC_50_ values were calculated using non-linear regression analysis in GraphPad Prism. Data are represented as mean ± SD, from at least three independent experiments.

|  | **Gem-miR-15a**<br>**(nM)** | **5-FU**<br>**(nM)** | **Fold**<br>**enhancement** |
| --- | --- | --- | --- |
| **Spheroid**<br>**SW620** | 9.75 ± 0.89 | 15,370 ± 2.3 | ~1576 |
| **CRC670** | 12.28 ± 0.72 | 8,004 ± 0.4 | ~652 |
| **CRC726** | 6.17 ± 0.87 | 1,300 ± 0.05 | ~211 |
| **HCRC4** | 13.43 ± 0.46 | 37,200 ± 23.3 | ~2770 |

### Gem-miR-15a inhibits metastatic tumor growth *in vivo* without detectable toxicity

A metastatic mouse model was established to investigate the therapeutic potential of Gem-miR-15a in CRC *in vivo*. Tumor xenografts were established by injecting luciferase expressing HCT116 cells (HCT116 + Luc) in NOD-SCID mice via tail-vein, followed by treatment with either vehicle control, Gem-miR-15a and Gem (Figure 7A). Tumor progression was monitored by bioluminescence imaging, which showed a significant reduction in the rate of tumor growth in both the Gem-miR-15a (4mg/kg) and Gem treated groups (50mg/kg), compared to the vehicle control (Two-way ANOVA, treatment x time effect **p=0.0038). Representative bioluminescence images demonstrating the reduced luciferase signal in the treated group as compared to control are shown (Figure 7B). Additionally, body mass was monitored throughout the study, to account for potential acute toxicity. No significant changes in body weight were observed, and the body mass remained in normal limits (<15% decrease in mass), indicating the absence of treatment-associated acute toxicity (Figure 7C). Collectively, these data indicate the therapeutic efficacy of Gem-miR-15a *in vivo*, thereby underlining its translational potential in CRC.

**Figure 7.**
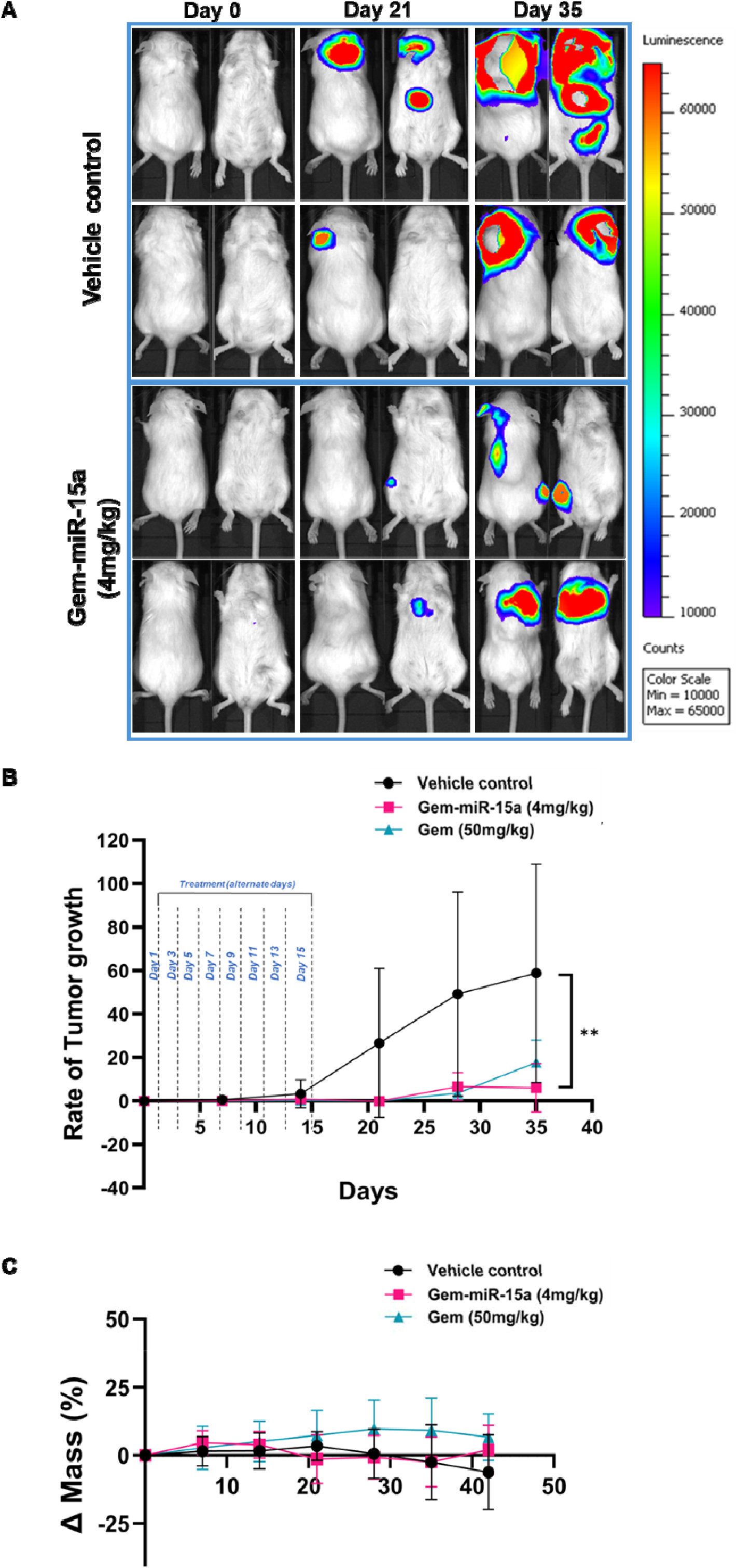
Gem-miR-15a suppresses metastatic tumor progression *in vivo* without detectable toxicity. (A) Representative *in vivo* bioluminescence (IVIS) images (dorsal and ventral views) of NOD/SCID mice bearing luciferase-expressing HCT116 metastatic tumors in vehicle control and Gem-miR-15a treated groups at indicated time points (Day 0, Day 21, Day 35). Progressive increase in signal intensity is observed in the vehicle group, whereas reduced tumor burden is evident in the Gem-miR-15a treated mice. (B) Quantification of rate of tumor progression over time based on bioluminescence signal intensity. Vehicle-treated mice exhibit a significantly higher rate of tumor growth compared to Gem-miR-15a (4 mg/kg, q2d x 8) and gemcitabine (Gem; 50 mg/kg, q3d x 4)-treated groups. (C) Body weight monitoring during treatment. No significant changes in body weight were observed across treatment groups. Tumor burden was monitored by bioluminescence imaging (n=4). Data are represented as mean ± SD. Statistical analyses were performed using two-way ANOVA with mixed effects. *p<0.05, **p<0.01, ***p<0.001, ****p<0.0001.

## Discussion

Resistance to 5-FU-based chemotherapy regimens poses a major challenge in the treatment of colorectal cancer (CRC) and significantly contributes to disease-related mortality (45). In this study a unique miRNA modification strategy was developed to improve stability and deliverability by introducing Gem into the guide strand on miR-15a. Gemcitabine-modified miR-15a (Gem-miR-15a) is a potent miRNA-based therapeutic that efficiently inhibits proliferation, induces S-phase cell cycle arrest and apoptosis, while downregulating key oncogenic proteins in both parental and resistant CRC cells. More importantly, Gem-miR-15a overcomes multiple forms of resistance, showing activity in clinically relevant 3D and *in vivo* models. Collectively, the findings establish Gem-miR-15a as a promising multi-targeted therapeutic approach that addresses several key limitations of conventional therapies.

It is well established that reduced miRNA-15a expression in CRC correlates with poor patient prognosis and although native miRNA-15a has therapeutic potential, a major obstacle to miRNA-based therapeutics is their delivery (15, 30, 46). Unmodified miRNAs are incapable of crossing the plasma membrane and require a delivery vehicle, which is often accompanied by toxicities (46, 47). Early attempts at miRNA-based therapeutics (MRX34) were unsuccessful, encountering multiple setbacks during Phase 1 clinical trials (17).

The present study establishes Gem-miR-15a as an effective modified miRNA therapeutic that retains anti-tumor activity *in vitro*, without the need for a delivery vehicle, thereby overcoming one of the persistent bottlenecks in RNA-based therapeutics. Gem modification does not alter the biological activity of miR-15a but renders it more stable with enhanced potency. The modification preserves Watson-Crick base pairing, as the fluorine group at the 2’ carbon of the ribose ring of Gem does not interfere with hydrogen bonding between nucleobases. Additionally, the incorporation of fluorine enhances the lipophilicity of the molecule, potentially facilitating cellular uptake and delivery (48, 49). Prior work has demonstrated that the Gem-miR-15a mimic has significant therapeutic potential in pancreatic cancer and can sensitize the pancreatic cancer cells to Gem (34). In this study, Gem-miR-15a suppressed established and functionally relevant oncogenic miR-15a targets, such as WEE1, CHK1, YAP1, BMI1 and CCND1 in CRC cells (Figure 4). This is mechanistically significant, since these targets represent an interconnected survival network involved in multiple oncogenic processes, imparting survival and stem-like characteristics to the cancer cells. Although these targets have been individually validated as therapeutic candidates, single-agent approaches targeting WEE1 and CHK1 have demonstrated limited clinical success (50-53), likely due to compensatory pathway activation (54). In contrast, the pleiotropic nature of Gem-miR-15a enables simultaneous targeting of multiple oncogenic drivers, thereby limiting compensatory adaptation and resistance development (55).

The multi-targeted suppression was strongly reflected in the phenotypic data, where Gem-miR-15a exhibited low-nanomolar growth inhibition, substantially exceeding the potency of 5-FU and oxaliplatin, with IC_50_ values approximately 100-5000-fold lower across all CRC cell lines. Gem-miR-15a was also more potent than native miR-15a in HCT116 cells, indicating enhanced efficacy upon Gem incorporation (Supplementary Figure 1).

Mechanistically, Gem-miR-15a consistently induced S-phase cell cycle arrest accompanied by depletion of the G2-cell population, as shown by the significant reduction in the G2/S ratio. This is consistent with the known mechanism of Gem, which incorporates into DNA and causes premature chain termination, thereby disrupting replication. The observed increase in the expression of γH2AX post-treatment further validates the induction of replication - associated DNA damage (39, 56). Suppression of WEE1 and CHK1 further impairs the cell’s ability to survive replicative stress, potentially pushing them towards mitotic catastrophe (57). By downregulating these proteins, Gem-miR-15a disables DNA damage checkpoint responses, leading to the accumulation of irreparable DNA lesions and ultimately apoptotic cell death. The observed downregulation of CCNA2 and CCNB1 provides additional support for cell cycle disruption and accumulation of cells in the S-phase (58, 59). Thus, the combined effect of Gem-induced replication disruption and miR-15A-mediated suppression of cell cycle checkpoint proteins synergistically drives robust cytotoxicity across all CRC cell lines (Figure 2A,4).

Consistent with these findings, robust apoptosis induction was observed following Gem-miR-15a treatment, evidenced by a significant increase in Annexin V-positive cells in both parental as well as resistant cells, further supported by an increased expression of cleaved PARP, cleaved Caspase-7, and pro-apoptotic BAK (Figure 2B).

RNA-seq analysis further supported the phenotypic observations and provided a better understanding of the potential mechanism of Gem-miR-15a in overcoming chemoresistance. Pathway enrichment analysis revealed modulation of key oncogenic and stress response pathways, including apoptosis, p53 signaling, PI3K-Akt, mTOR, Hippo, autophagy and inflammatory response signaling, all of which are critically implicated in CRC progression and therapy resistance (Figure 5). Dysregulation of PI3K-Akt and mTOR pathways has been extensively associated with enhanced tumor survival, metabolic adaptation and resistance to fluoropyrimidine-based therapies, while aberrant Hippo signaling and *YAP1* activation, have been linked to stemness, metastasis and poor prognosis in CRC (60-62). Similarly, autophagy has been reported to support tumor cell survival under chemotherapeutic stress, thereby contributing to resistance (63, 64).

Notably, Gem-miR-15a treatment resulted in the coordinated suppression of several resistance-associated pathways, while simultaneously activating stress and apoptotic pathways. At the gene level, downregulation of *AKT3* and *MAP2K6*-associated signaling is consistent with prior studies implicating PI3K-Akt and MAPK pathways in mediating resistance to both 5-FU and oxaliplatin in CRC (65-68). In parallel, upregulation of key apoptotic mediators such as *APAF1, BAX* and *FAS* along with DNA damage-associated genes including *RAD51C, DDB2, XPC*, and *EXO1*, supports activation of replication stress-induced cell death pathways (69, 70). The induction of these genes reflects an attempt to repair treatment-induced DNA damage, with concurrent apoptotic signaling indicating DNA damage accumulation beyond repair capacity, ultimately leading to cell death (71).

The observed downregulation of *AURKA*, a mitotic kinase frequently overexpressed in CRC, is particularly notable, as Aurora kinase-A inhibition has been shown to enhance sensitivity to 5-FU and impair tumor cell proliferation (72, 73). Additionally, activation of canonical p53 signaling and downstream targets such as *CDKN1A* (p21) is consistent with prior reports demonstrating the critical role of p21 in mediating response to both 5-FU and oxaliplatin in CRC (74).

Importantly, resistance-reversal analysis demonstrated that Gem-miR-15a not only suppresses key survival pathways such as mTOR, HIF-1 and Hippo signaling but also restores apoptotic signaling through the induction of genes such as *CASP10*, which has been linked to chemotherapy-induced apoptotic regulation (60, 61, 75, 76). Collectively, these findings indicate that Gem-miR-15a exerts its anti-tumor effects through coordinated inhibition of pro-survival signaling and activation of DNA damage-associated stress response pathways, effectively shifting resistant cells from a pro-survival toward a pro-apoptotic state (Figure 5, Supplementary Figure 5).

A major highlight of this study is the ability of Gem-miR-15a to overcome drug-resistance, maintaining potent activity in 5-FU-resistant and oxaliplatin-resistant CRC cells (Figure 3). These findings not only highlight the intrinsic potency of Gem-miR-15a but also underscores its ability to restore therapeutic sensitivity in tumors that have acquired resistance to standard chemotherapies.

In the 5-FU-resistant model, untreated resistant cells showed an increased level of thymidylate synthase (TS), a well-established mechanism of fluoropyrimidine resistance, which was reduced upon Gem-miR-15a treatment (Figure 5). This suggests a plausible mechanism by which Gem-miR-15a reverses 5-FU resistance. Additionally, broad suppression of cell cycle checkpoint regulators and survival pathways likely aid in the re-sensitization at a system’s level, suggesting that the resistance-reversal phenotype is probably multifactorial (45).

The ability of Gem-miR-15a to overcome oxaliplatin resistance is noteworthy in light of prior literature suggesting that Gem can bypass oxaliplatin resistance in CRC through inhibition of Akt and p38/MAPK pathways (40). Consistent with this, transcriptomic analysis revealed downregulation of key signaling mediators including *AKT3, MAP2K6*, along with the suppression of mTOR pathway, which has been implicated in Gem-mediated anti-tumor activity (Figure 5). Given the central role of Akt-mTOR axis in CRC resistance, its coordinated inhibition likely contributes to the observed reversal of oxaliplatin resistance. Future studies should further dissect the molecular mechanism underlying oxaliplatin resistance and its reversal.

The synergy observed between Gem-miR-15a and oxaliplatin in parental HCT116 cells, reflected by both the Bliss synergy score of 14 and a high combination sensitivity score, highlights a clinically relevant translational strategy (Figure 6). Rather than viewing Gem-miR-15a solely as a replacement strategy, the data supports its potential use in rational combinations to enhance response and overcome resistance.

Given the central role of genetic determinants in modulating therapeutic response in CRC, *TP53* mutations account for approximately 60% of CRC cases and are frequently associated with resistance and poor clinical outcome (77, 78). Interestingly, Gem-miR-15a remained highly effective in both HCT116 (p53^-/-^, p21^+/+^) and HCT116 (p53^+/+^, p21^-/-^) cells, demonstrating that its anti-tumor activity is largely independent of the p53-p21 axis (Figure 1,2). This p53-independence is particularly encouraging given the extensive heterogeneity of *TP53* alterations in CRC. While the RNA-seq and protein data indicate stress-induced activation of p53 pathway in p53-proficient cells, the retained efficacy in knockout models suggests that Gem-miR-15a engages additional p53-independent mechanisms of cytotoxicity, suggesting that p53 may augment, but is not a necessity for the efficacy of Gem-miR-15a (Figure 5, Supplementary Figure 3). Future mechanistic studies should define how Gem-miR-15a mediates cell death in the absence of p53. The ability to bypass p53 dependency, in contrast to p53-restoration strategies explored in CRC, may confer broader therapeutic advantage across genetically diverse tumor types.

In this study, Gem-miR-15a showed strong efficacy not only in the spheroid-forming CRC cells, but also across multiple patient-derived organoids, from both primary and metastatic origin further demonstrating its clinical relevance (Figure 6). Notably, tumor organoids preserve the genetic heterogeneity and phenotypic characteristics of the original tumors, making them physiologically superior models for therapeutic response assessment compared to conventional cell lines (79, 80).

In the metastatic CRC model, both Gem-miR-15a and Gem reduced the rate of tumor growth relative to the vehicle control, at doses of 4 mg/kg and 50 mg/kg, respectively. Gem-miR-15a achieved anti-tumor activity at a substantially lower dose than free Gem, with no significant body weight loss or overt toxicity observed. This dosing advantage is particularly relevant because Gem has not succeeded as a standard CRC therapy; however, evidence supports its role as a salvage therapy in heavily pretreated or refractory CRC cases (81, 82). Additionally, low dose Gem has been shown to act as an immunomodulator by selectively depleting regulatory T cells in the tumor microenvironment (42, 43). These findings raise the possibility that chemical reprogramming, rather than conventional administration of the free drug, may unlock therapeutic value of Gem in CRC.

Future studies should evaluate the pharmacokinetics and immune-modulatory effects of Gem-miR-15a, particularly in immunocompetent or syngeneic mice. Given the modulation of inflammatory pathways observed in RNA-seq analysis (Figure 5), it would be interesting to determine whether Gem-miR-15a influences anti-tumor immunity.

Advances in oligonucleotide therapeutics are increasingly supporting the feasibility of chemically modified RNA drugs, especially when they can reduce dependence on toxic delivery vehicles. Notably, a Phase 1 clinical trial of 5-FU-miR-15a (CR1-02) in relapsed/ refractory Acute myeloid leukemia (AML) patients confirmed its safety and tolerability with early signs of biological activity (83). These findings collectively support Gem-miR-15a as a promising multimodal therapeutic candidate for CRC. The molecule simultaneously addresses three major challenges: the poor deliverability of native miRNAs, pathway redundancies underlying resistance to single-target therapies, and acquired resistance to standard chemotherapy, thereby positioning Gem-miR-15a as a novel strategy for overcoming therapeutic resistance in CRC.

## Conclusions

In summary, Gem-miR-15a represents a potent, multi-targeted therapeutic that effectively suppresses CRC growth by inducing replicative stress, cell cycle arrest and apoptosis. By simultaneously targeting multiple oncogenic drivers, Gem-miR-15a efficiently overcomes resistance to 5-FU-based chemotherapies, with robust activity in clinically relevant 3D organoid models and *in vivo*, at substantially reduced doses without observable toxicity. Collectively, these findings establish Gem-miR-15a as a promising therapeutic candidate with strong potential for overcoming chemoresistance and improving treatment outcomes.

## List of abbreviations

Gem-miR-15a: gemcitabine modified miRNA-15a
5-FU: 5-fluorouracil
CRC: Colorectal cancer
miRNA: MicroRNA
miR-15a: MicroRNA-15a
Gem: Gemcitabine
PDAC: Pancreatic ductal adenocarcinoma

## Declarations

### Ethics approval and consent to participate

Patient-derived organoids were obtained from the Stony Brook Medicine Biobank under an approved institutional review board (IRB) protocol.

All animal experiments were approved by the Institutional Animal Care and Use Committee (IACUC) of Stony Brook University and were conducted in accordance with institutional guidelines and applicable regulations for animal care and use.

### Consent for publication

Not applicable

### Availability of data and materials

RNA sequencing data generated in this study will be deposited in GEO and will be available as of the date of publication. All other data supporting the findings of this study are available with the corresponding author, upon request. Any additional information required to reanalyze the data reported in this paper is available from the corresponding author, upon request.

### Authors’ contributions

AO and JJ conceptualized the study; AO and AP performed the data curation, investigation and analysis; MC, AO and RD performed the RNA-seq and bioinformatics analysis; AO wrote and edited the original manuscript; AO, AP and JJ made critical revisions; JJ supervised the study and acquired funding and resources.

## Acknowledgements

We thank all members of Dr. Ju’s laboratory for the valuable insights, constructive feedback and continued support throughout this project. The graphical abstract was created using BioRender.

---

**Supplementary figure 1.**
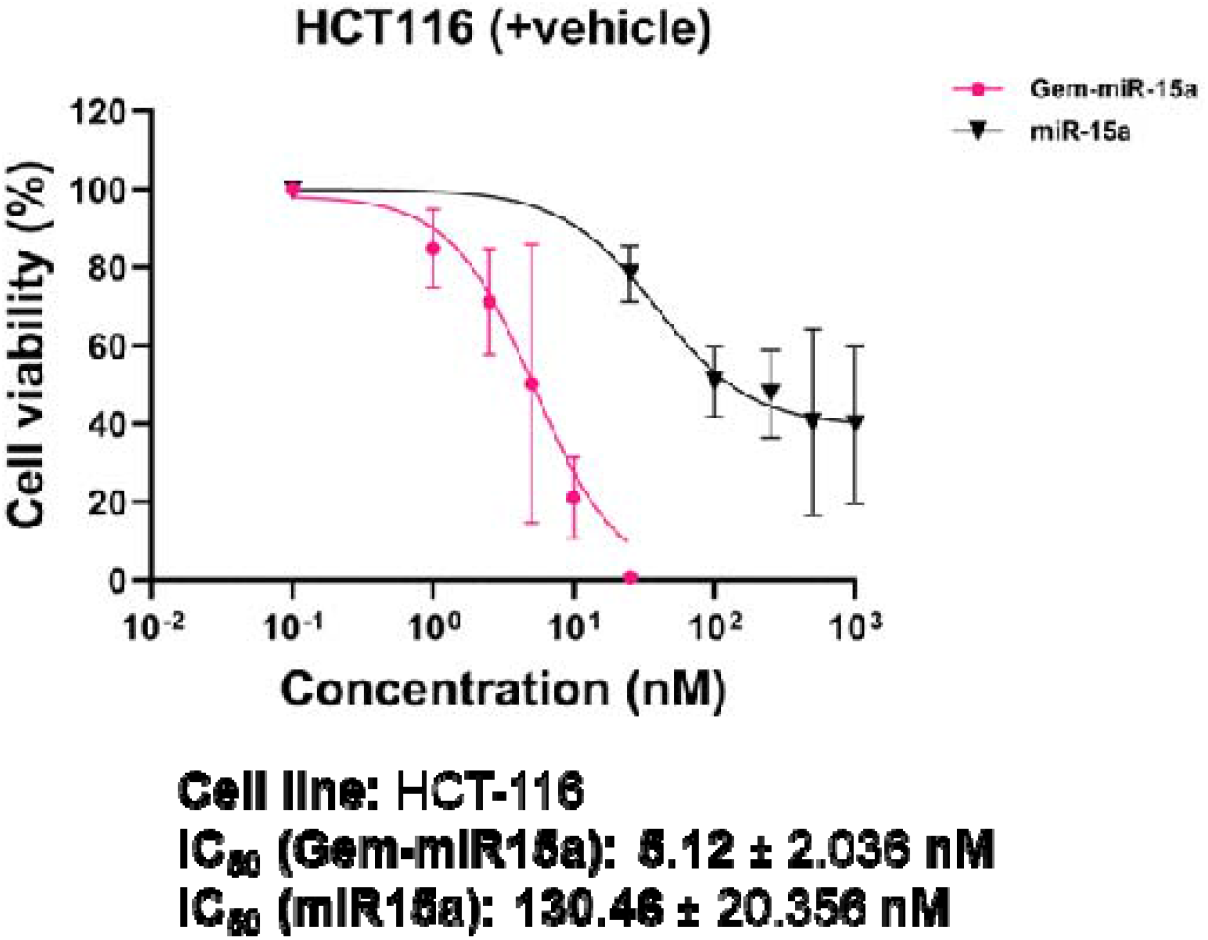
Gem-miR-15a exhibits enhanced potency compared to native miR-15a, related ic Figure 1. Dose response analysis comparing the effect of native miR-15a and Gem-miR-15a on cell viability in the presence of vehicle control. Cells were treated with increasing concentrations of each mimic and the viability was assessed 6-days post-transfection. Gem-miR-15a demonstrated significantly enhanced potency with an IC_50_ value of 5.12 nM compared to 130.46 nM for native miR-15a.

**Supplementary figure 2.**
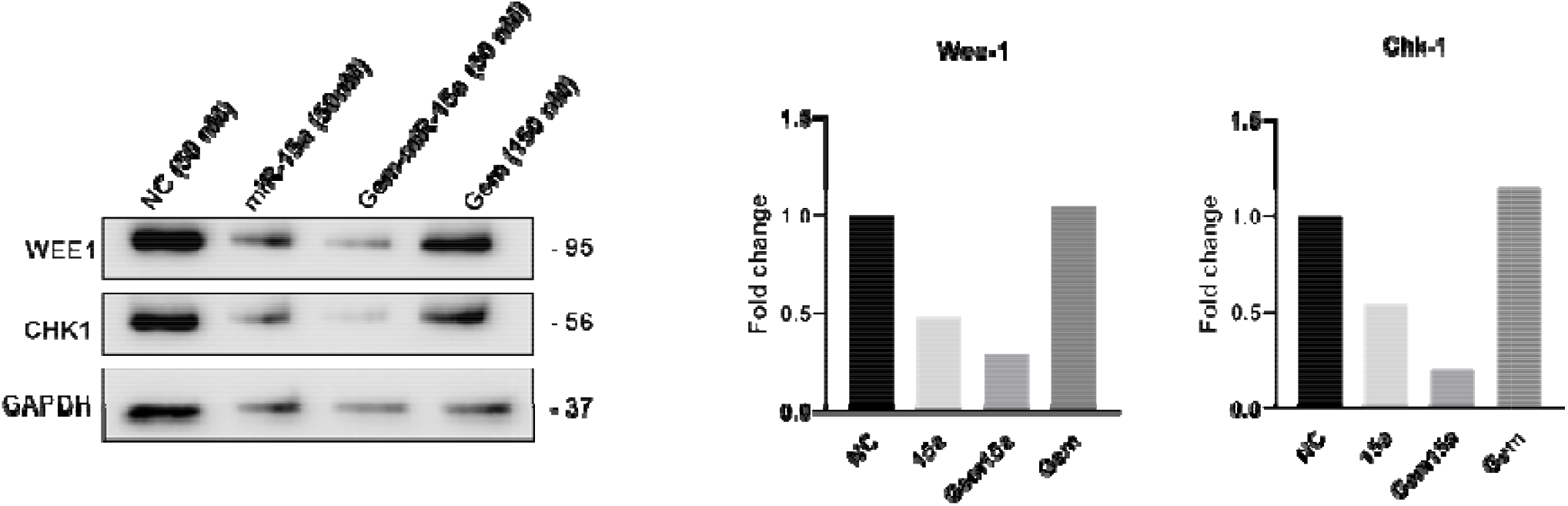
Gem-miR-15a suppresses target proteins In 5-FU-resistant CRC cells, related to Figure 4. Western blot analysis of 5-FU Res -ICT116 cells treated with NC, miR-15a, Gem-miR-15a or gemoltabine (Gem), showing suppression of established miR-15a or gemoltabine (Gem) showing suppression of established miR-15a targets WEE1 and CHK1.

**Supplementary figure 3.**
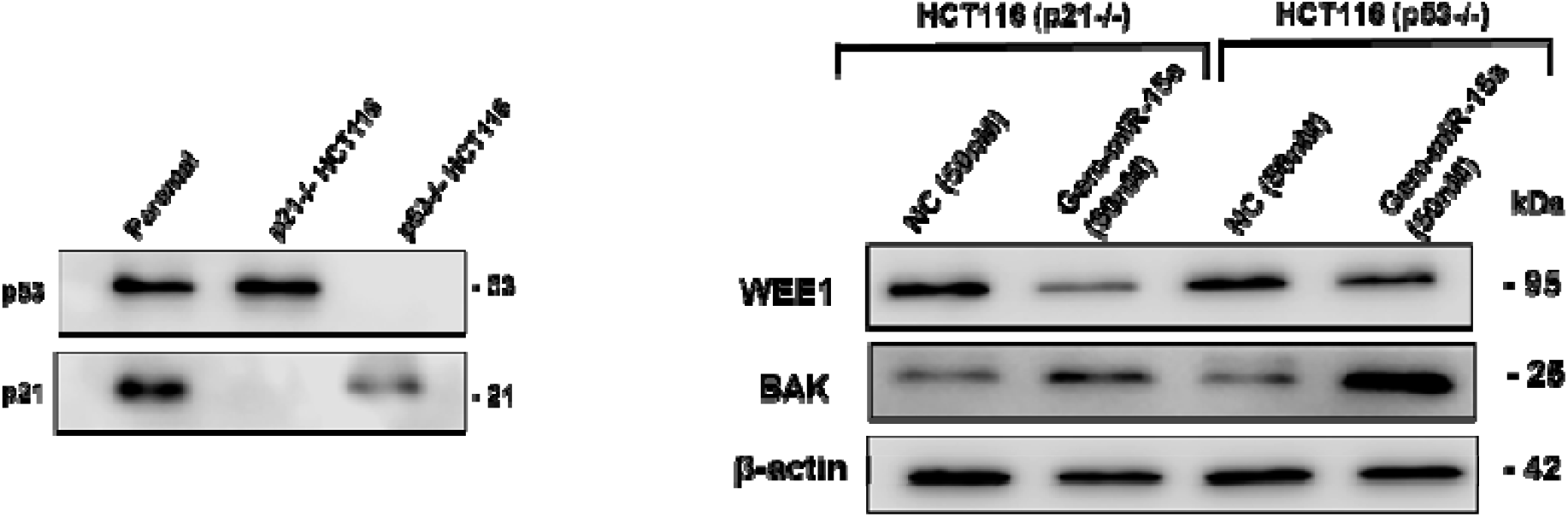
Validation of p53 and p21 knockout and downstream effects of Gem-miR-15a, related to Figures 2. Western blot analysis confirming knockout of p53 and p21 In HCT116 p53’ and p21’ cells. In addition, treatment with Gem-miR-15a results In downregulation of WEE1 and Induction of pro-apoptotic BAK, compared to NC, Indicating activation of apoptotic signaling Independent of p53/p21 status.

**Supplementary figure 4.**
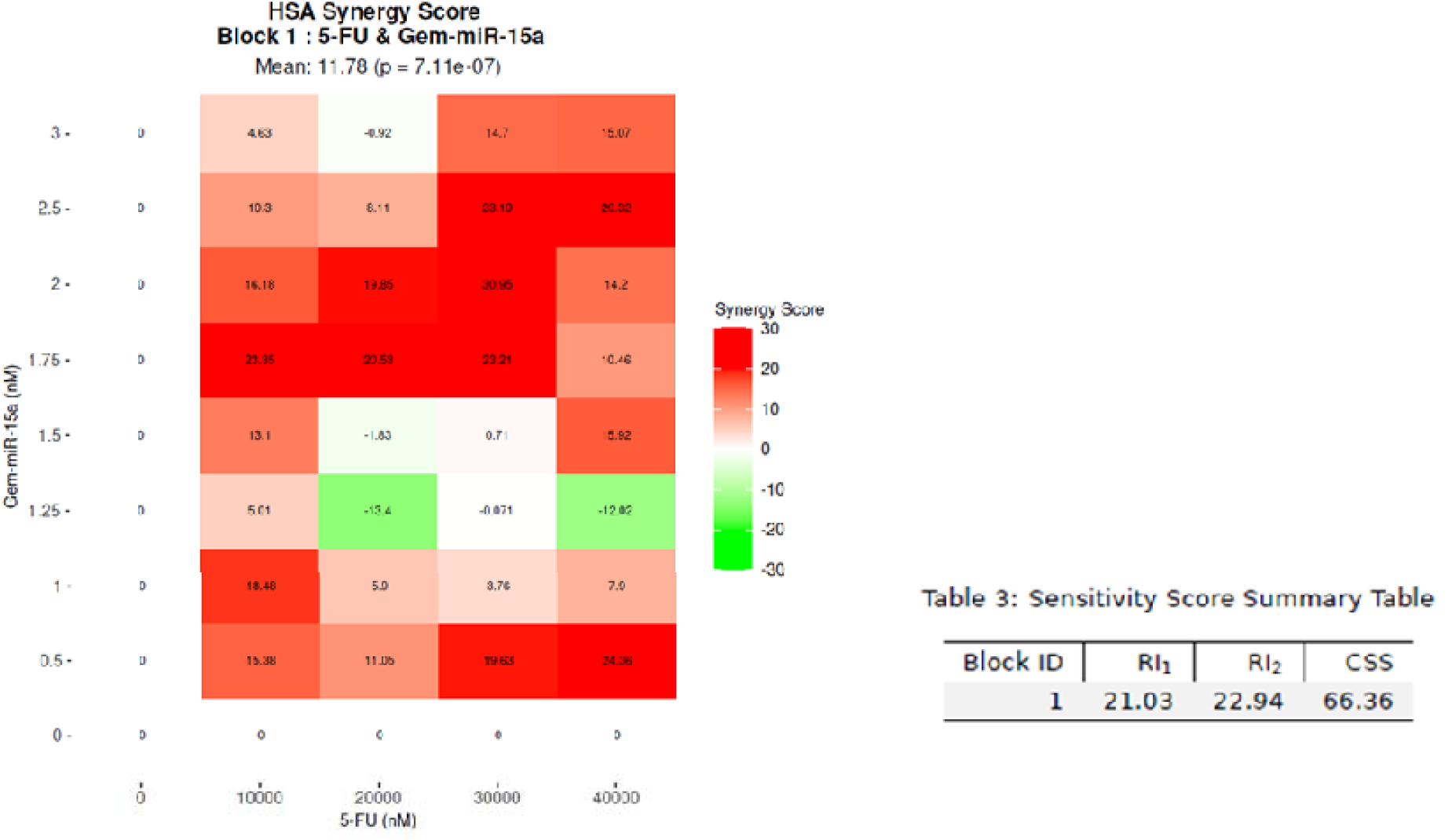
Gem-miR-15a exhibits synergistic Interaction with 5-FU in resistant CRC cells, related to Figures 3. Synergy analysis of Gam-miR 15a in combination with 5-FU in 5-FU Res HCT116. Heatmap representation of drug interaction demonstrates a synergistic effect, with a Highest single agent (HSA) synergy score of 11.78 and a combination sensitivity score (CSS) of 66.36. Data are derived from dose-response matrix analysis using SynergyFinder+.

**Supplementary figure 5.**
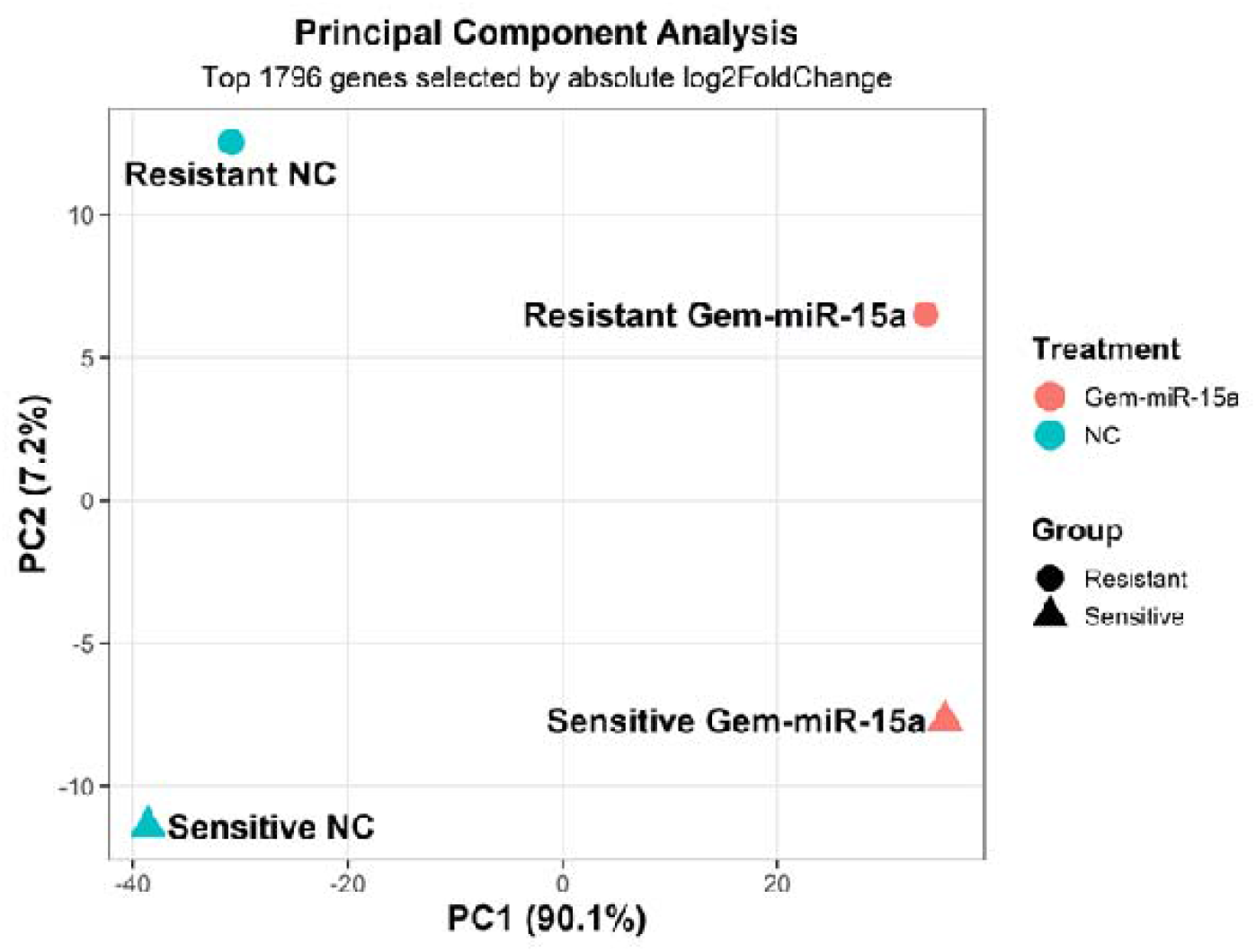
Gem-miR-15a treatment shifts transcriptomic profiles of resistant CRC cells toward parental state, related to Figures 5. Principal component analysis (PCA) of gene expression profiles in parental and 5-FU Res - ICT116 cells under untreated and Gem-miR-15a treated conditions. Untreated parental and resistant cells form distinct clusters, Indicating divergent transcriptomic states. Following Gem-miR-15a treatment, both parental and resistant samples shift toward a common cluster, with reduced separation between groups, suggesting partial convergence of resistant cells toward a parental-like transcriptional profile.

